# A novel RRM activity orchestrates ribosome homeostasis

**DOI:** 10.64898/2026.04.18.719249

**Authors:** James A. Saba, Phoebe E. White, A. Maxwell Burroughs, L. Aravind, Rachel Green

## Abstract

Ribosome concentration varies across cell type, state, and disease, yet the mechanism by which mammalian cells sense ribosome levels to regulate ribosome synthesis remains unknown. Previous studies implicate LARP1 in binding ribosomes and, separately, in repressing mRNAs containing TOP motifs (TOPs), which encode all ribosomal proteins as well as several translation factors. Here, we show that ribosome binding is the essential licensing event for LARP1-mediated TOP repression. Phylogenetic analysis reveals that LARP1’s ribosome binding region is part of a highly conserved RNA recognition motif (RRM) which is predicted to directly interact with its TOP-binding HEAT repeat domain. Ribosome binding activates the HEAT repeats to bind and repress TOPs, thereby repressing TOP translation and ribosome synthesis. Mutants which uncouple the coordinated functions of these domains constitutively repress TOPs and compromise cell fitness. These data reveal an unprecedented ribosome sensing function of LARP1, orchestrated through the unique coordinating role of its RRM, unveiling the mechanism by which mammalian cells tune ribosome synthesis to demand across diverse contexts.

## Introduction

The ribosome is an essential macromolecular complex for all life. Eukaryotic ribosomes comprise a 40S and 60S ribosomal subunit composed in total of four ribosomal RNAs (rRNAs) and ∼80 ribosomal proteins (r-proteins), and are expressed in the range of hundreds of thousands to millions of copies per cell^1,2^. The process of ribosome synthesis is both indispensable and energetically costly, and thus highly regulated^1–3^. While the ribosome itself represents an impressive proportion of cellular biomass^4,5^, ribosome concentration varies across cell type, state, and disease process^6–9^. A fundamental molecular mechanism that determines how mammalian cells sense ribosome levels to enact precise regulation of ribosome synthesis has not been established.

La-related protein 1 (LARP1) was long known to encode a C-terminal HEAT repeat domain which binds and represses Terminal OligoPyrimidine motif-containing mRNAs (TOPs)^10–12^, which encode all ∼80 r-proteins as well as several translation factors^13^. We and others identified a new region within LARP1, the ribosome binding region (RBR), which directly binds the mRNA channel of the 40S ribosomal subunit^14,15^. Given these distinct binding activities, we speculated that LARP1 might couple them to repress TOPs (and thus ribosome synthesis) when ribosome levels exceed cellular demand. Although non-translating ribosomes were recruited to LARP1’s RBR by any stress that increased free 40S subunits, a mechanism directly implicating ribosome binding in the pathway of TOP repression remained elusive^14^. With the advent of AlphaFold3, more advanced predictions of inter– and intramolecular protein contacts permit more precise design of mutants to probe this potential cellular function^16^.

Here, we show through AlphaFold3 prediction, evolutionary conservation, and *in vitro* and *in cellulo* biochemistry, that LARP1 functions as a *bona fide* sensor of ribosomes that directly regulates ribosome homeostasis. Our analysis suggests that LARP1’s RBR and HEAT repeat domains interact via a previously unrecognized RNA recognition motif (RRM) domain, found in other La-related proteins (LARPs)^17–20^ but thought to be absent in LARP1^21,22^. We demonstrate that interactions predicted to hold the RRM and HEAT repeats together are incompatible with 40S subunit binding to LARP1, supporting a model in which 40S binding to LARP1’s RBR activates the C-terminal HEAT repeats to bind and repress TOPs. Consistent with this model, 40S binding to LARP1 is required for LARP1-mediated TOP repression in cells, indicating that this binding is the critical licensing event for repression. Together, these interactions define an elegant sensing mechanism by which LARP1 samples free 40S subunits to shut down ribosome synthesis when ribosome levels exceed cellular demand.

## Results

### Evolutionary conservation of the region spanning the La-wHTH and HEAT repeat domains

LARP1 contains an N-terminal La winged helix-turn-helix domain (La-wHTH)^22,23^ and a C-terminal DM15 region containing a HEAT repeat domain^10,11^ (Figure 1A). Between these domains, prior work from our group identified the 40S-binding RBR as a series of conserved helical segments connected by disordered regions, consistent with Alphafold2^24^ (AF2)-based modeling (Figure 1A)^14^. The release of AlphaFold3 (AF3)^16^ prompted us to revisit the region between the La-wHTH and HEAT repeats. Surprisingly, AF3 modeled a core globular region featuring a 4-stranded beta sheet (*β*S) with two ɑ-helices stacking on one face of the sheet (Figure 1A and Supplemental Figures S1A-B). Specifically, residues corresponding to the core structured elements of this globular domain are I643-Q648 (*β*1), A673-W691 (ɑ1), S798-V803 (*β*2), V832-M836 (*β*3), P875-N883 (ɑ2), and G884-V889 (*β*4) (Figure 1A and Supplemental Figure S1C). The first helix (ɑ1) corresponds to RBR helix 1 (H1), which binds the 40S mRNA channel^14,15^, and the second helix (ɑ2) and fourth beta strand (*β*4) lie immediately upstream of the HEAT repeats^10,11^.

**Figure 1.**
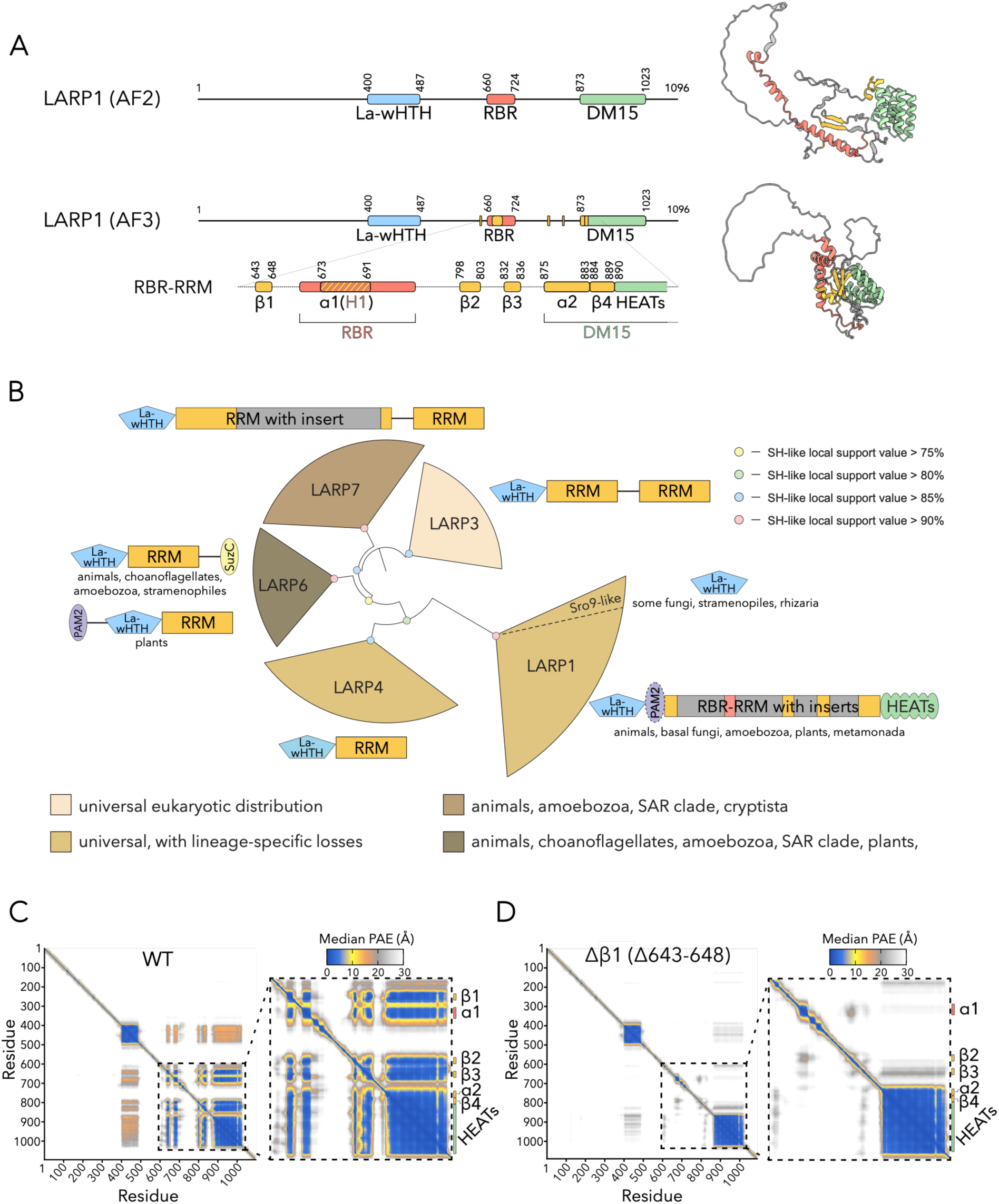
The LARP1 RBR-RRM is predicted to coordinate the 40S-binding ɑ1 helix to the TOP-binding HEAT repeat domain. **A)** Comparison of AlphaFold2 (AF2) and AlphaFold3 (AF3) predicted structures of LARP1. Top: primary structure (to scale) of LARP1 with corresponding AF2 prediction. Bottom: primary structure (to scale) of LARP1 with corresponding AF3 prediction (single model, representative of ten models; Supplemental Figure S1A). In the primary structure, the RRM domain (yellow) and RBR domain (red) are shown, with the RRM ɑ1 helix corresponding to helix 1 (H1) of the RBR depicted with yellow and red dashes. For AF models, the region spanning the RBR-RRM (red and yellow) to HEAT repeats (green) (amino acids 640-999) is shown for clarity. **B)** Phylogenetic tree depicting relationships between LARP homologs, based on multiple sequence alignment spanning the La-wHTH and RRM domains (Supplemental Figure S3, Supplemental Table 1). LARP families on the tree are color-coded by phylogenetic distribution. Node color indicates Shimodaira–Hasegawa (SH)-like local support values. Representative domain architectures for each family are provided adjacent to clades. Gray boxes inside RRM domains represent low-complexity insert regions. Genbank accessions for representative architectures are as follows: *H. sapiens* LARP1, EAW61633.1; *S. cerevisiae* Sro9p, DAA07447.1; *H. sapiens* LARP3, EAX11258.1; *H. sapiens* LARP4A, EAW58140.1; *H. sapiens* LARP6, ANM63599.1; *A. thaliana* LARP6, ANM63599.1; *H. sapiens* LARP7, AAH66945.1. **C)** Median predicted aligned error (PAE) plot for WT-LARP1 from ten AF3 predictions (five models generated from each of two independent seeds). Zoom-in of the region spanning the RBR-RRM and HEAT repeats is shown to the right. Low PAE values (blue) demonstrate that the entire RBR-RRM and HEAT repeats are predicted to associate with each other. **D)** Identical to (**C**) except for Δ*β*1-LARP1 (Δ643-648). High PAE values (grey to white) demonstrate that the *β*S is not predicted to form and that the 40S-binding ɑ1 helix is not predicted to associate with the HEAT repeats.

Structural homology searches initiated with the region between the La-wHTH and HEAT repeats consistently recovered diverse representatives of the RRM superfamily^25^. For example, human LARP1 (Genbank: NP_291029.2) recovers the second RRM domain of the human U2 snRNP complex component HTATSF1 (PDB: 6Y53 chain q, TM-score: 0.67) and the first RRM repeat of the human Msx2-interacting protein (PDB: 4P6Q chain A, *Z*-score: 4.5); *B. floridae* LARP1 (GenBank: XP_035681167.1) recovers the first RRM domain of *D. melanogaster* SNF protein (PDB: 6F4I, TM-score: 0.60) and the first RRM repeat of human Msx2-interacting protein (PDB: 4P6Q chain A, *Z*-score: 4.6); *F. alba* LARP1 (Genbank: KCV70036.1) recovers the first RRM domain of human MARF1 protein (PDB: 2DIU chain A, TM-score: 0.67) and the yeast Nab3 RRM domain (PDB: 2XNQ chain A, TM-score: 0.60). We named this previously unrecognized RRM the RBR-RRM, as it includes the previously identified RBR within it (Figure 1A). While ɑ2 and *β*4 lie within the previously defined DM15 region^10,21^, we refer to them as part of the RBR-RRM and, for clarity, refer to the C-terminal HEAT repeats of the DM15 region specifically as “HEAT repeats”.

Despite the confirmed presence of RRM domains in other LARP protein families (La/LARP3-^17^, LARP4-^18^, LARP6-^19^, and LARP7-like^20^ (Supplemental Figure S2A)), support for an RRM domain in LARP1 has been debated^21,22^. LARP1’s RRM reported here is atypically long at the primary sequence level (Supplemental Figure S2B) and contains highly divergent loops of low complexity (Supplemental Figure S2C). These long, diversified loops between individual secondary structural elements result in “twilight zone” scores for the above structural homology searches^26,27^. We suspect that they have hampered detection of LARP1’s true RRM domain to this point.

To assess conservation of the RBR-RRM domain across eukaryotes, we initiated sequence homology searches spanning the start of *β*1 to the end of *β*4 as queries. We recovered RBR-RRM domains in LARP1 orthologs broadly spanning the eukaryotic tree of life, including in animals, choanoflagellates, filasterea, certain fungi, amoebozoans, plants, and the excavate Metamonada lineage (Figure 1B, Supplemental Figure S3, and Supplemental Table 1). Our analysis, building on prior analyses studying the evolutionary origin of the La-wHTH and HEAT repeats^10,21,28^, indicates for the first time that a three-domain LARP1 architecture (La-wHTH, RBR-RRM, and HEAT repeat domain) was present in the last eukaryotic common ancestor (Figure 1B). It likely emerged via duplication of the core La-wHTH and RRM domains, followed by acquisition of the C-terminal HEAT repeat domain and accretion of long disordered loops within the RBR-RRM. This trio of domains was subsequently vertically inherited across much of the eukaryotic tree of life, consistent with selective pressure for maintaining these domains. Beyond the classic three-domain LARP1 homologs, fungi possess the Sro9-SLF1-like family, which lacks both the RBR-RRM and HEAT repeats (Figure 1B and Supplemental Table 1); a similar situation is observed in short LARP1-like homologs from Rhizaria and Stramenopile lineages (Figure 1B and Supplemental Table 1). Some additional LARP1-like proteins across eukaryotes appear to lack one or more of the core domains; these could reflect true loss of these domains or genome assembly and prediction errors (Supplemental Table 1).

### LARP1 structural prediction

We next analyzed predicted structural contacts of the newly-identified RBR-RRM. AF3 models of full-length LARP1 revealed expected folding for the individual La-wHTH, RBR-RRM ɑ1, and HEAT repeats as previously observed in solved structures of these domains^11,14,22^ (Supplemental Figures S4A-D). Focusing on the RBR-RRM, contacts between the central *β*S and ɑ1 are predicted to form the core globular structure (Supplemental Figure S5); ɑ2 is positioned immediately N-terminal to *β*4 but is not otherwise predicted to interact with the remainder of the RBR-RRM beyond hydrophobic contacts typical of a folded structure. Strikingly, extensive contacts are also predicted between the RBR-RRM and the HEAT repeats, modeling a compound RBR-RRM/HEAT repeat scaffold. High-confidence interactions include predicted hydrogen bonds (H-bonds) between D637 (upstream of *β*1) with N928/K930 (HEAT); S798/F800 (*β*2) with N680 (ɑ1); R799 (*β*2) with E687 (ɑ1); Y801/P802 (*β*2) with R917 (HEAT); E829 (upstream of *β*3) with R668 (ɑ1); S830 (upstream of *β*3) with Q910 (HEAT); M836 (*β*3) with W691 (ɑ1); S838 (*β*3) with L690 (ɑ1); and Q887 (*β*4) with Y686 (ɑ1) as well as a predicted pi-stacking interaction between W834 (*β*3) and F922 (HEAT) (Supplemental Figures S5B-K). Importantly, AF3 reveals very low predicted aligned error (PAE) values between amino acids corresponding to the RBR-RRM and the HEAT repeat domain, supporting both the folding of the core RBR-RRM and predicted interactions with the HEAT repeats (Figure 1C).

We next introduced *in silico* deletions intended to disrupt the core *β*S of the RBR-RRM and observed what happens to AF3-predicted interactions between ɑ1 and the HEAT repeats. As expected, AF3 does not predict an intact *β*S when residues corresponding to either the central *β*1 (Δ643-648) or *β*3 (Δ832-836) strand is removed (Supplemental Figures S6A-B and S7A-B). More interestingly, in both *in silico* deletions the RBR-RRM ɑ1 and HEAT repeats (although still predicted to adopt their individual secondary structures) are no longer predicted to associate with each other, as indicated by PAE values (Figure 1D and Supplemental Figure S7C). Based on these predictions, we hypothesize that the core *β*S of the RBR-RRM holds the ɑ1 helix and C-terminal HEAT repeats in a coupled array.

Taken together, LARP1’s 40S-binding helix (ɑ1) and TOP-binding HEAT repeats are predicted to be physically connected in the form of a compound scaffold with the core *β*S of the RBR-RRM. Along with their evolutionary co-inheritance, these predictions provide a potential basis for an integrated function of these modules.

### Disrupting the predicted RBR-RRM/HEAT scaffold induces constitutive 40S binding and TOP repression

Our *in silico* predictions suggest that the 40S-binding ɑ1 helix and the TOP-binding HEAT repeats interact as components of a compound scaffold with the core *β*S of the RBR-RRM. One possible implication is that these modules are mutually repressed through interactions with the core *β*S of the RBR-RRM. To probe this possibility, we mutated the predicted core *β*S of the RBR-RRM. We designed three mutants for *in cellulo* analysis: 1) *β*1-P (L644P/T647P), a double proline substitution in the *β*1 strand, 2) *β*1-del (Δ643-647), a deletion of the *β*1 strand, and 3) *β*3-del (Δ832-836), a deletion of the *β*3 strand. These mutants all target central strands of the *β*S and, we predict, would disrupt it entirely, potentially activating the HEAT repeats and ɑ1 helix to bind TOPs and 40S subunits, respectively (schematic in Figure 2A).

**Figure 2.**
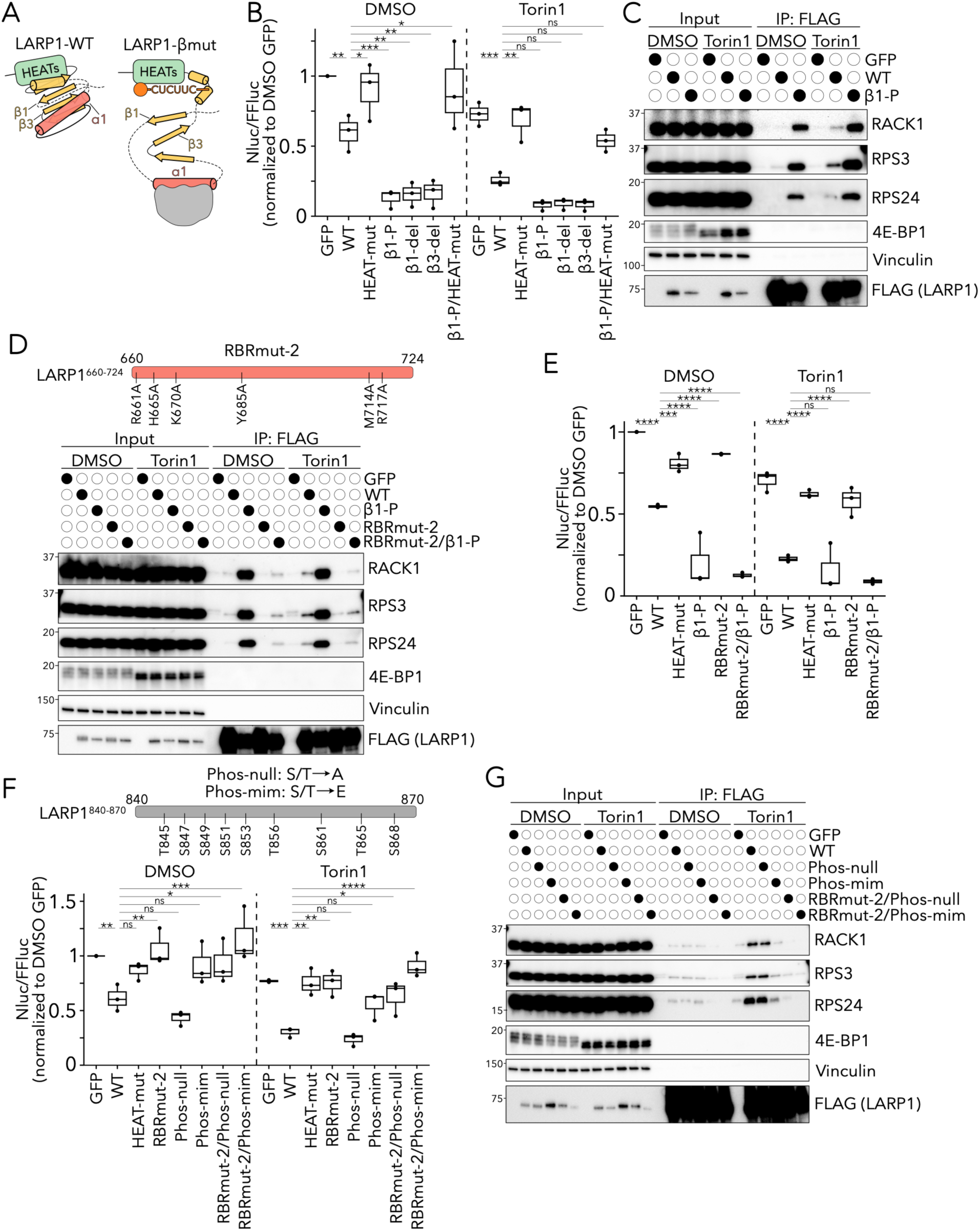
Disrupting the predicted RBR-RRM/HEAT scaffold activates 40S binding and TOP repression. **A)** Hypothesized model depicting structural and functional predictions for WT (left) and *β*-mut (right) LARP1. In WT LARP1, the RBR-RRM/HEAT scaffold is predicted to be functionally repressed, with minimal 40S binding to the ɑ1 helix and TOP binding to the HEAT repeats. In *β*-mut LARP1, the central *β*S of the RBR-RRM is predicted to be disrupted, activating the ɑ1 helix and HEAT repeats to bind 40S subunits and TOPs, respectively. The RBR-RRM (yellow), ɑ1 helix (red), HEAT repeat domain (green), 40S subunit (grey), and TOP mRNA (orange) are depicted. The La-wHTH domain is omitted. **B)** Expression of the TOP (RPL39-nLuc-PEST) reporter^14^. LARP1-KO cells were transfected with plasmids expressing the indicated LARP1 construct (or GFP) and treated with DMSO or 300 nM Torin1 for 1 hour. NLuc expression was normalized to FFLuc expressed on the same plasmid. Data points represent the mean of three technical replicates from three independent biological replicates. Whiskers represent minima and maxima; bounds of box represent the quartiles (25th and 75th percentile); center line represents the median. Values are normalized to the GFP-transfected, DMSO-treated sample. Two-way ANOVA with Dunnett’s correction for multiple comparisons; n.s. not significant, * p ≤ 0.05, ** p ≤ 0.01, *** p ≤ 0.001, **** p ≤ 0.0001. **C)** Immunoblots of input (0.05%) or elution (15%) following immunoprecipitation of FLAG-tagged GFP or the indicated FLAG-tagged LARP1^574-1096^ construct. Constructs were expressed in LARP1-KO cells treated with DMSO or Torin1 (300 nM) for 1 hour. Data is representative of at least two independent biological replicates. **D)** Top: Schematic depicting the LARP1 RBR (residues 660-724) with the indicated point mutations used in RBRmut-2 LARP1. Bottom: Same setup as (**C**) for the indicated LARP1 constructs. **E)** Same setup as (**B**) for the indicated LARP1 constructs. **F)** Top: Schematic depicting the 840-870 amino acid region of LARP1 with mutated residues in Phos-null LARP1 (Ala substitutions) or Phos-mim LARP1 (Glu substitutions). Bottom: Same setup as (**B**) for the indicated LARP1 constructs. **G)** Same setup as (**C**) for the indicated LARP1 constructs.

To probe the extent to which these mutants bind and repress TOPs, we used a previously validated reporter in which a TOP 5’UTR is expressed upstream of a nanoluciferase-PEST (nLuc-PEST) open reading frame^14^. The nLuc-PEST protein is rapidly degraded and therefore reports acutely on the translation status of the mRNA. As a control, we used a non-TOP reporter with the same design except preceded by a non-TOP 5’UTR. Wild-type (WT) LARP1 and the *β*S mutants were expressed at similar levels to endogenous LARP1 in LARP1-KO cells (Supplemental Figure S8A). As seen previously^14^, expression of WT-LARP1 induces moderate repression of the TOP reporter, which is enhanced by Torin1 (an mTOR inhibitor^29^) and abolished by a previously validated HEAT repeat mutant (R917E/Y960A^10,11^; HEAT-mut) (Figure 2B). WT-LARP1 has little effect on translation of the non-TOP reporter (Supplemental Figure S8B). When we express any mutant predicted to disrupt the *β*S, we observe strongly increased TOP repression compared to WT (Figure 2B); this repression is essentially constitutive, as it no longer responds to Torin1. This repression further depends on the function of the HEAT repeat domain, as a *β*1-P/HEAT double mutant does not repress the TOP mRNA (Figure 2B). Surprisingly, we also observe mild repression of the non-TOP reporter with all three *β*S mutants (Supplemental Figure S8B), likely reflecting enhanced binding of the HEAT repeats to non-TOPs despite low affinity for these alternative motifs^10,11^. Together, these data suggest that disrupting the proposed interaction network of the RBR-RRM and the HEAT repeats renders the HEAT domain constitutively active.

We next probed ribosome binding using a co-immunoprecipitation (co-IP) approach. Notably, the La-wHTH domain and PAM2 motif of LARP1 bind polyA tails and polyA-binding protein, respectively^22,30^. To avoid co-IPing mRNAs via their polyA tails (and thus non-specifically co-IPing ribosomes on mRNAs), we used an N-terminally truncated version of LARP1 in which the La-wHTH and PAM2 regions are removed (LARP1^574–1096^)^12^. We expressed FLAG-tagged truncated LARP1 (Supplemental Figure S8C), IPed FLAG, and immunoblotted for co-IPed 40S r-proteins which reflect 40S subunits bound to LARP1. Compared to WT, *β*1-P mutant LARP1 co-IPs substantially more 40S subunits in both untreated and Torin1-treated conditions (Figure 2C and Supplemental Figure S8D), suggesting that disrupting the proposed RBR-RRM/HEAT interaction renders ɑ1 more available for 40S binding.

These results support a model in which the RBR-RRM functionally represses the TOP-binding HEAT repeat domain and the 40S-binding ɑ1 helix. Disruption of this predicted core architecture appears to license each element to engage in its respective binding activity.

### Ribosome binding is required for TOP repression in cells

We previously proposed the model that LARP1’s RBR might function as a sensor of 40S subunits, preventing the synthesis of new ribosomes when non-translating 40S subunits are in excess^14^. To test this model and guided by AF2, we generated a multi-point mutant of LARP1 which was unable to bind ribosomes (RBRmut-1; Supplemental Figure S9A)^14^. While initially excited by the hypothesis that RBRmut-1 would be unable to repress TOPs, we were surprised to find that it repressed TOPs to an even greater extent than WT^14^.

Informed by our new insights from AF3, evolutionary analysis, and the *β*S mutants, we wondered whether RBRmut-1 introduced mutations in the ɑ1 helix that, in addition to disrupting ribosome binding, inadvertently disrupt the predicted RBR-RRM/HEAT scaffold. Indeed, according to AF3, three residues mutated in RBRmut-1 are predicted to interact with the central *β*S as part of the core globular structure of the RBR-RRM: R668 H-bonds with E829 (upstream of *β*3; Supplemental Figure S5F), Y686 H-bonds with Q887 (*β*4; Supplemental Figure S5J), and W691 H-bonds with M836 (*β*3; Supplemental Figure S5H). We find that mutating just these three residues (RYW-mut) induces TOP repression equivalent to that observed for RBRmut-1 (Supplemental Figures S9A-C). We propose that this repression reflects disruption of the predicted RBR-RRM architecture in a manner analogous to the *β*S mutants.

Having clarified the shortcomings of our prior mutant, we revisited the 40S sensing model. Notably, based on the cryo-EM structure^14^, 40S binding to the ɑ1 helix of LARP1’s RBR-RRM would directly clash with the AF3-predicted folding of the RBR-RRM/HEAT scaffold (Supplemental Figures S10A-C). This comparison suggests a model wherein the 40S subunit, in binding to the ɑ1 helix, could disrupt the predicted RBR-RRM/HEAT scaffold in a manner analogous to our *β*S mutants. While this broad structural rearrangement is inferred from the AF3 structural prediction of LARP1, it would provide a potential molecular basis for LARP1 to directly sense 40S ribosomal subunits and transduce this signal to activation of the HEAT repeats. Consistent with this model, we note that *in silico* deletion of the ɑ1 helix (Δ673-691), modeling a major disruption of this region as might occur upon 40S binding, is predicted by AF3 to disrupt the RBR-RRM and disengage the HEAT repeats (Supplemental Figures S11A-C).

Using AF3 and the RBR-40S structure as a guide, we generated a new mutant that is predicted to disrupt the interaction with the 40S subunit without impinging on predicted interactions within the RBR-RRM (RBRmut-2; Figure 2D and Supplemental Figure S9A). Mutation of six residues rendered RBRmut-2 deficient for binding 40S subunits (Figure 2D and Supplemental Figures S9D-E). Remarkably, when introduced into our reporter assay, RBRmut-2 fails to repress the TOP reporter even in response to Torin1; this mutant effectively phenocopies the HEAT repeat mutant, implicating 40S binding as a required step in LARP1-mediated TOP repression (Figure 2E). The RBRmut-2 defect is bypassed with a RBRmut-2/*β*1-P double mutant, which activates the HEAT repeats even in the absence of 40S binding (Figures 2D-E).

These data establish 40S binding as an essential step in LARP1-mediated TOP repression. They further suggest that 40S binding functions mechanistically by activating the HEAT repeats, as the requirement for a 40S subunit can be overcome by mutations that constitutively activate the HEAT repeats. Finally, these data provide strong support for the model that LARP1’s RBR-RRM ɑ1 helix acts as a sensor for free ribosomes.

### LARP1 phosphorylation antagonizes 40S binding

Current models implicate mTOR as a major regulator of LARP1 and TOP translation^31^. LARP1 directly interacts with mTOR and is dephosphorylated upon mTOR inhibition^9,32^, modulating TOP binding and repression^12,33^. Having observed that 40S binding is required for LARP1 to repress TOPs, we next interrogated interactions between LARP1 phosphorylation and 40S binding in the context of TOP repression.

To probe these interactions, we selected nine LARP1 serine or threonine residues that are phosphorylated by mTOR to regulate TOP translation^33^; we mutated these residues to alanines (Phospho-null; “Phos-null”) or glutamates (Phospho-mimetic; “Phos-mim”) (schematic in Figure 2F). Consistent with predictions, Phos-null LARP1 increased repression of the TOP reporter relative to WT, whereas Phos-mim LARP1 diminished the repression (Figure 2F). Preventing 40S binding by adding the RBRmut-2 point mutations on top of the Phos-null or Phos-mim mutant derepresses the TOP reporter to LARP1-KO levels (Figure 2F). These data provide orthogonal support to our earlier observations by demonstrating that 40S binding is required for TOP repression independent of LARP1 phosphorylation status. When we measured ribosome binding using these mutants, Phos-null LARP1 displayed equivalent 40S binding compared to WT whereas Phos-mim LARP1 displayed impaired 40S binding, specifically under conditions of mTOR inhibition (Figure 2G and Supplemental Figure S9F). These data indicate that sustained LARP1 phosphorylation antagonizes 40S binding. We speculate that, while de-phosphorylated LARP1 may be more amenable to 40S binding than phosphorylated LARP1, when cells are translationally active and there are few free 40S subunits, we fail to capture these binding differences.

Taken together, these results demonstrate that LARP1 phosphorylation antagonizes binding of the RBR-RRM ɑ1 helix to available 40S subunits. We propose that phosphate groups on LARP1 repel the negatively-charged surface of the 40S subunit or, alternatively, stabilize the predicted RBR-RRM/HEAT interaction, thus limiting 40S access to the ɑ1 helix.

### LARP1 couples 40S binding and TOP binding in vitro

We next developed an *in vitro* reconstituted system to more fully explore the biochemical parameters by which 40S binding to LARP1’s RBR-RRM mediates TOP binding and repression by the HEAT repeats. We purified truncated FLAG-tagged LARP1^574–1096^ containing the RBR-RRM and HEAT repeats (Supplemental Figure S12A)^12^, 40S ribosomal subunits (Supplemental Figure S12B), and ^32^P-labeled capped TOP or non-TOP oligos (Supplemental Figure S12C). We co-incubated LARP1^574–1096^ with equivalent amounts of either TOP or non-TOP oligo and increasing amounts of 40S subunits and performed a FLAG immunoprecipitation to measure oligo binding to LARP1 as a function of 40S subunit concentration (Experimental setup in Figure 3A). In this assay, we find that LARP1^574-1096^ binds the TOP oligo more strongly than the non-TOP oligo (Figure 3B and Supplemental Figure S12D), consistent with prior reports^10,11^. Moreover, the addition of 40S subunits dramatically increases LARP1^574-1096^ binding to the TOP oligo in a dose-responsive manner (Figure 3B and Supplemental Figure S12D). In contrast, there was no effect of 40S subunits on LARP1^574-1096^ binding to the non-TOP oligo (Figure 3B and Supplemental Figure S12D). These data demonstrate that the 40S subunit is sufficient to drive TOP, but not non-TOP, binding to LARP1 in a minimal system.

**Figure 3.**
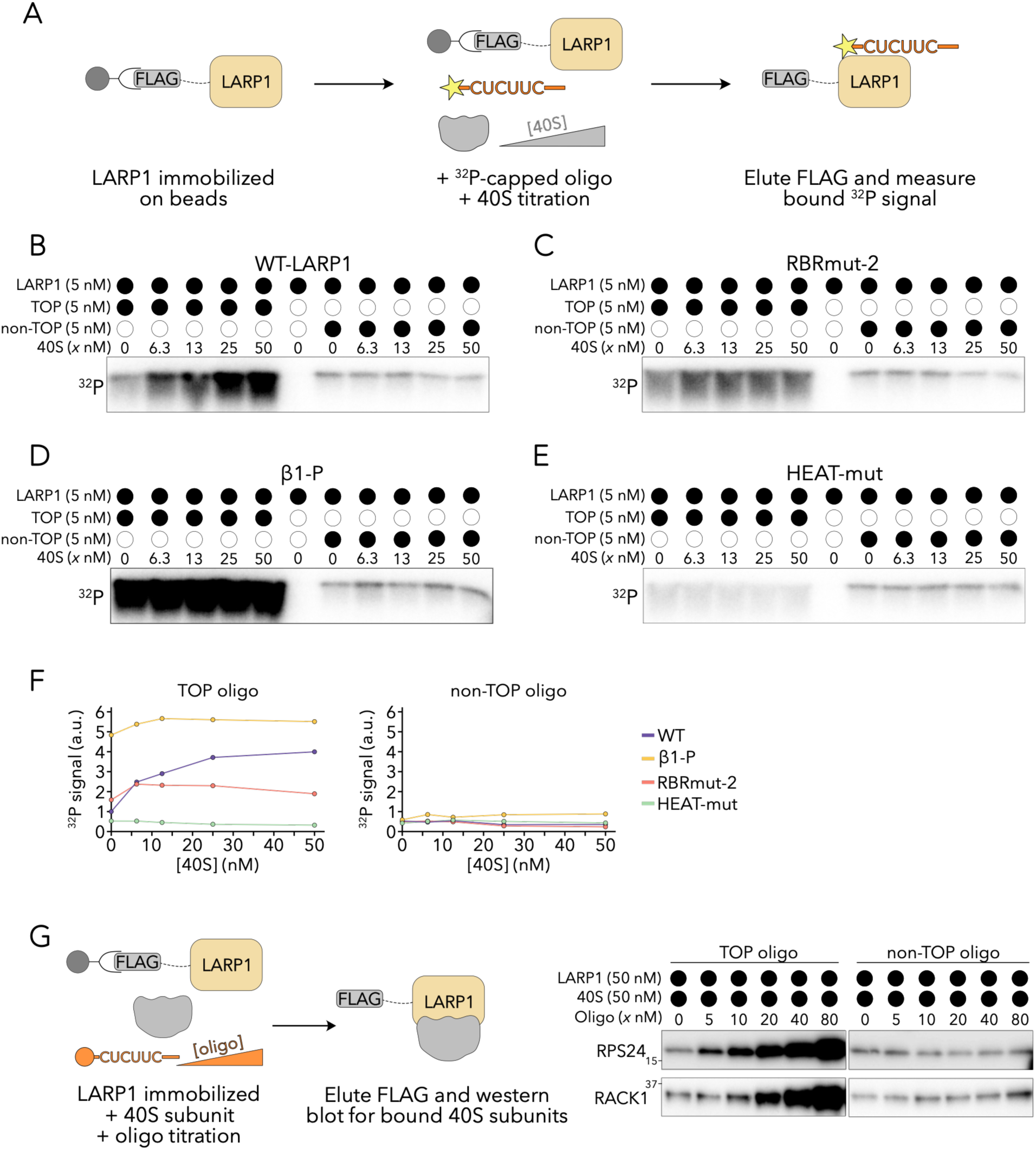
LARP1 couples 40S binding and TOP binding *in vitro*. **A)** Schematic depicting experimental setup for Figures 3B-E. Purified, FLAG-tagged LARP1^574-1096^ was bound to anti-FLAG beads and then co-incubated with a constant amount of ^32^P-capped TOP or non-TOP oligonucleotide and varying amounts of 40S subunits. The beads were washed and eluted, followed by measurement of ^32^P signal on a gel. A stronger ^32^P signal indicates that more oligonucleotide was bound by LARP1 during the incubation. **B-E)** Experimental results for setup shown in (**A**) using WT-LARP1 (**B**), RBRmut-2 LARP1 (**C**), *β*1-P LARP1 (**D**), or HEAT-mut LARP1 (**E**). Panels (**B-E**) were performed in a single experiment; levels of bound oligonucleotide can be directly compared between panels. Results are representative of at least two independent experiments. **F)** Quantification of ^32^P signal intensity in (**B**-**E**) (a.u. arbitrary units). Values are normalized to WT LARP1 incubated without 40S subunits. **G)** Left: Experimental setup. Purified, FLAG-tagged LARP1^574-1096^ was bound to anti-FLAG beads and then co-incubated with a constant amount of 40S subunits and varying amounts of capped TOP or non-TOP oligonucleotide. The beads were washed and eluted, followed by measurement of bound 40S ribosomal proteins by immunoblot. A stronger signal for bound 40S ribosomal proteins indicates that more 40S subunits were bound by LARP1 during the incubation. Right: Experimental results. Results are representative of three independent experiments.

We also tested the 40S-stimulatory effect with various LARP1 mutants. Importantly, 40S stimulation of TOP binding is completely lost with LARP1^574-1096^ variants carrying mutations in the ɑ1 helix, the *β*S, or the HEAT repeat domain (Figures 3C-F and Supplemental Figures S12E-H). Specifically, RBRmut-2 LARP1, incapable of binding ribosomes through its ɑ1 helix, reveals intermediate TOP binding but is unresponsive to 40S subunits (Figure 3C and Supplemental Figure S12E). The *β*1-P mutant, which constitutively activates the HEAT repeats, exhibits sustained strong binding to the TOP oligo independent of 40S subunits (Figure 3D and Supplemental Figures S12F). Finally, the HEAT repeat mutant, incapable of binding TOPs, demonstrates constitutive weak binding (Figure 3E and Supplemental Figure S12G); this last observation suggests that mutations in the HEAT repeat that prevent TOP binding also disrupt signal transduction though the RBR-RRM/HEAT scaffold, consistent with a predicted interaction between one of the residues mutated in the HEAT mutant and *β*2 of the RBR-RRM (Supplemental Figure S5E). The cumulative results of these experiments are quantified in Figure 3F and Supplemental Figure S12H. These data provide direct biochemical evidence for the 40S subunit driving TOP binding, which requires signal transduction through the ɑ1 helix, *β*S, and HEAT repeat domains.

While we found that 40S binding to LARP1 drives TOP binding, we also wondered whether TOP binding might similarly stimulate 40S binding. To this end, we co-incubated LARP1^574-1096^ and 40S subunits with increasing amounts of either TOP or non-TOP oligo, followed by FLAG immunoprecipitation (Experimental setup in Figure 3G). Using immunoblotting to quantify r-proteins and assess 40S binding to LARP1, we find that the non-TOP oligo has no effect on 40S binding, while addition of the TOP oligo strongly increases 40S binding in a dose-responsive manner (Figure 3G). We further note that *in silico* deletions predicted to disrupt the RBR-RRM induce an AF3-predicted refolding of the ɑ2 helix and *β*4 strand, with *β*4 adopting a helical conformation contiguous with the first HEAT repeat (Supplemental Figure S12I). This conformation mirrors that observed in the solved crystal structure of the HEAT repeats bound to a TOP oligo ^11^. This alternative conformation provides an initial hypothesis for how the 40S binding signal could be biophysically transduced to the HEAT repeats and, *vice versa*, how TOP binding could be biophysically transduced to the RBR-RRM.

Collectively, these data demonstrate that either 40S binding to the ɑ1 helix or TOP binding to the HEAT repeat domain of LARP1 is sufficient to drive binding of the other component. These findings are consistent with a cooperative mode of 40S and TOP binding^34^ and provide direct biochemical support for a coordinated role of LARP1’s RBR-RRM and HEAT repeats in executing TOP repression.

### Disrupting the predicted RBR-RRM/HEAT scaffold perturbs endogenous TOPs and cell fitness

Having established the coordinated role of the RBR-RRM and HEAT repeats in regulating TOPs in reporter-based assays and *in vitro*, we next examined their coordination in regulating endogenous TOPs. We performed RT-qPCR of sucrose gradient fractions from LARP1-KO cells overexpressing WT, *β*1-P, or RBRmut-2 LARP1 (Supplemental Figures S13A-B). Following sucrose gradient fractionation, we split the gradient into two populations corresponding to sub-polysomes (mRNAs with zero or one ribosome loaded) or polysomes (mRNAs with multiple ribosomes loaded). In cells expressing WT-LARP1, TOPs are distributed between sub-polysomes and polysomes, consistent with previous reports of TOPs displaying a biphasic distribution^14,35–37^ (Figure 4A and Supplemental Figure S13C). In cells expressing *β*1-P LARP1, TOPs are almost entirely shifted to the sub-polysome population, indicating strong translation repression (Figure 4A and Supplemental Figure S13C). In contrast, in cells expressing RBRmut-2 LARP1, TOPs are shifted into the polysome population, indicating translation activation (Figure 4A and Supplemental Figure S13C). Non-TOPs sediment almost entirely in the polysome population in a manner unaffected by LARP1 variant expression (Figure 4B and Supplemental Figure S13C). These data demonstrate that the coordinated activity of LARP1’s RBR-RRM and HEAT repeats is required for translation regulation of endogenous TOPs.

**Figure 4.**
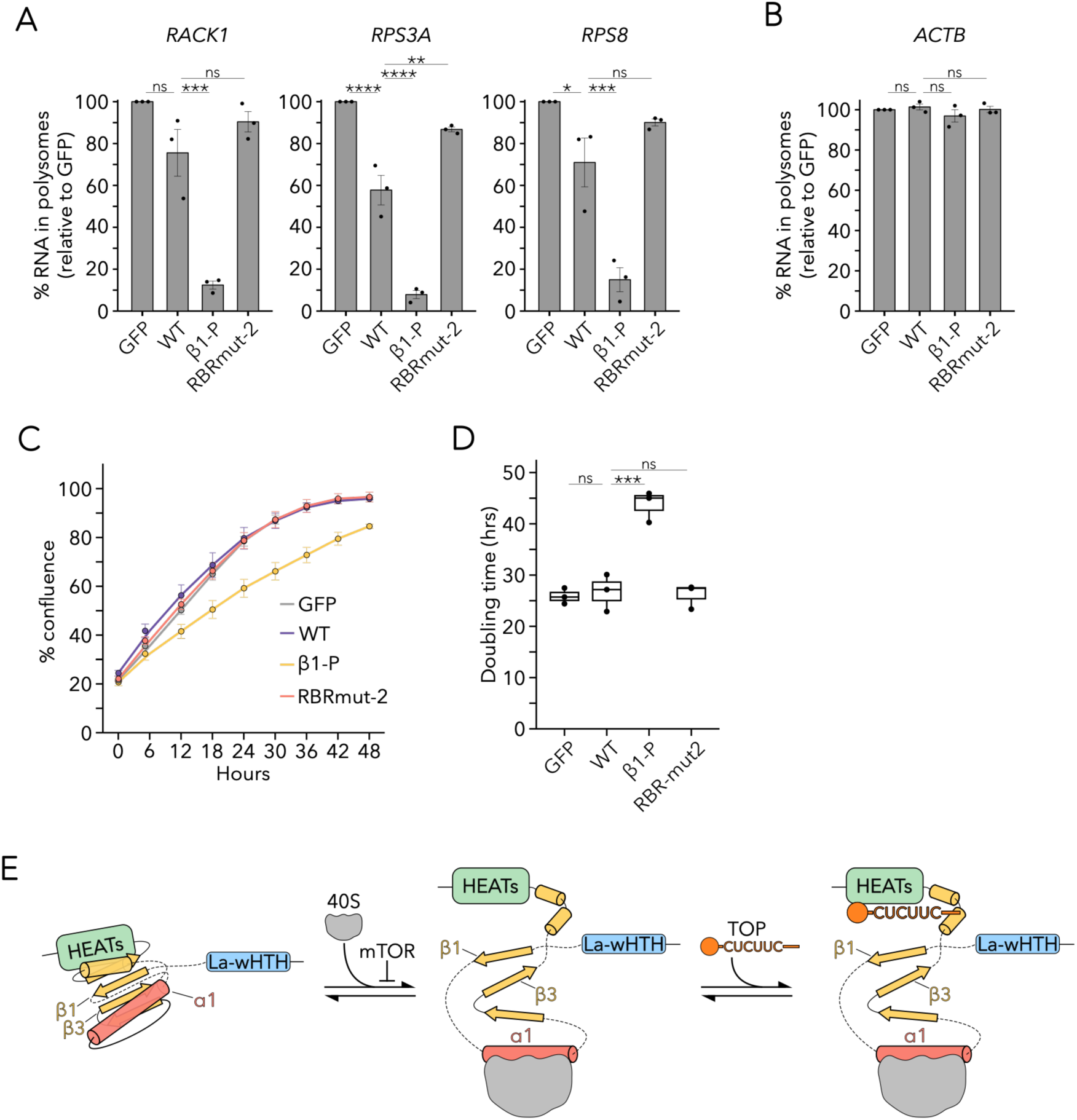
The predicted RBR-RRM/HEAT scaffold regulates endogenous TOPs. **A)** Percentage of TOP mRNAs in polysome fractions. LARP1-KO cells expressing the indicated LARP1 construct (or GFP) were fractionated over a sucrose gradient and qPCR for the indicated TOP was performed from pooled populations. Bars represent the mean of three independent biological replicates. Data points represent the mean of three technical replicates from three independent biological replicates and error bars represent the SEM. Values are normalized to the GFP-transfected sample. One-way ANOVA with Dunnett’s correction for multiple comparisons; n.s. not significant, * p ≤ 0.05, ** p ≤ 0.01, *** p ≤ 0.001, **** p ≤ 0.0001. **B)** Same setup as (**A**) for the indicated non-TOP. **C)** Growth curves over 48 hours of LARP1-KO cells transfected with the indicated LARP1 construct (or GFP) at endogenous levels. Data points represent the mean of three independent biological replicates each with three technical replicates. Error bars represent ± one standard deviation across biological replicates. **D)** Quantification of doubling time from growth curves presented in **C**). Data points represent the mean of three technical replicates from three independent biological replicates. Whiskers represent minima and maxima; bounds of box represent the quartiles (25th and 75th percentile); center line represents the median. One-way ANOVA with Dunnett’s correction for multiple comparisons; n.s. not significant, *** p ≤ 0.001. **E)** Model: The LARP1 RBR-RRM functions as a ribosome-responsive regulator of TOP mRNAs. 40S subunit binding to the ɑ1 helix is predicted to disrupt the RBR-RRM/HEAT architecture, releasing the HEAT repeats to bind TOPs and repress their translation. Phosphorylation of LARP1 by mTOR antagonizes 40S binding. The La-wHTH domain (blue), RBR-RRM (yellow), ɑ1 helix (red), HEAT repeat domain (green), 40S subunit (grey), and TOP mRNA (orange) are depicted.

In addition to TOP repression, LARP1 has been shown to stabilize TOP mRNAs^32,36,38^. We thus tested whether LARP1’s RBR-RRM similarly coordinates LARP1-mediated TOP stabilization, using TOP steady state levels as a proxy for stability. To this end, we transfected LARP1-KO cells with WT, *β*1-P, or RBRmut-2 LARP1 for 48 hours and performed RT-qPCR to evaluate endogenous TOP levels. Introduction of WT-LARP1 into LARP1-KO cells modestly increases TOP steady state levels, and this effect is strongly increased by the *β*1-P mutant (Supplemental Figure S13D). In contrast, RBRmut-2 LARP1 fails to show any effect on TOP steady state levels (Supplemental Figure S13D). Together, these data argue that the coordinated activity of LARP1’s RBR-RRM and HEAT repeats is involved in the stabilization of endogenous TOPs.

Finally, we investigated how expression of *β*1-P or RBRmut-2 LARP1 affects cell proliferation. We tracked the growth of LARP1-KO cells transfected with WT, *β*1-P, or RBRmut-2 LARP1 for 48 hours. In this assay, the growth of LARP1-KO cells expressing the *β*1-P mutant was compromised, revealing a doubling time of ∼40 hours compared to ∼24 hours for LARP1-KO cells expressing either GFP or WT-LARP1 (Figures 4C and 4D). These data suggest that constitutive HEAT activation and consequent TOP repression is detrimental to cell growth. In contrast, the growth of cells expressing RBRmut-2 LARP1 was unperturbed, suggesting that constitutive activation of TOPs in nutrient-replete conditions is not problematic. A similar growth defect of the *β*1-P mutant was observed upon conditions of recovery from serum starvation (Supplemental Figure S13E). Taken together, these data establish that 40S binding to LARP1’s RBR-RRM regulates endogenous TOPs, and that proper regulation of the LARP1-driven 40S sensing network is essential for cell fitness.

## Discussion

Here, we establish that LARP1 is a *bona fide* sensor of ribosomes that directly regulates ribosome homeostasis. The previously identified 40S-binding helix of LARP1 is part of a divergent and previously unrecognized RRM domain (termed here the RBR-RRM), homologous to the RRM domain observed C-terminal to the La-wHTH throughout eukaryotic LARP families^17–20^. Our AF3-predictions and biochemical data suggest a normally inactive state of LARP1’s RBR-RRM in which the 40S-binding ɑ1 helix and C-terminal HEAT repeat domain directly interact with the central *β*S. Our data further support a cooperative mode of binding, wherein the 40S subunit activates the C-terminal HEAT repeats to bind the 5’-cap of TOPs and repress their translation. While the exact biophysical mechanism by which 40S binding activates the HEAT repeats is not known, extensive clashes with the AF3-predicted structure of the RBR/RRM-HEAT scaffold suggest that 40S binding could drive conformational unfolding of the RBR-RRM and release of the HEAT repeats to bind TOPs. Ultimately, this coordinated activity allows cells to directly sense non-translating 40S subunits and repress the synthesis of ribosomal proteins as a function of 40S availability (Model Figure 4E). Considering early studies revealing feedback from ribosomal proteins to rRNA^39^ and the requirement of ribosomal proteins for induced folding of nascent rRNA^40–43^, we propose that this mechanism serves a broad and fundamental role in the regulation of ribosome synthesis as a whole.

The RRM domain is one of the most abundant and functionally diverse domains in all eukarya, distributed across varied nucleic acid-binding roles^44^. While RRMs are thought to adopt a stable structure, induced fit binding to nucleic acid targets has been reported^45,46^. To our knowledge, our discoveries reveal the RBR-RRM of LARP1 as the first example in which a folded RRM might be structurally remodeled upon binding its target (the 40S subunit) to promote the “activity” of the protein. In the case of LARP1, the acquisition of long disordered loops between secondary structural elements of the RBR-RRM could provide flexibility that facilitates rearrangement of the otherwise stable globular core upon 40S binding. Despite variability in sequence, these disordered loops are an ancient and persistent feature of LARP1 across eukaryotes, suggesting that functional coupling between 40S binding and HEAT repeat activation might be a widely conserved feature. Similarly, an insert in the RRM of LARP7 suggests that it could undergo an analogous mechanism of activation relevant to its role in transcription^47^.

Many cellular stresses impinge upon translation initiation, often through the action of mTOR or the integrated stress response^48^, leading to ribosome runoff and consequent binding of free 40S subunits to LARP1^14^. Moreover, different cellular regimes (e.g. stem cell maintenance or differentiation^6^, cancers^49^ and disease states^50^) have different requirements for ribosome abundance; here, too, non-translating 40S subunits should serve to determine ribosome homeostasis through LARP1 binding. Multiple factors – SERBP1 (a ribosome hibernation factor^51,52^), RIOK3 (a 40S degradation factor^53^), and NSP1 (a SARS-CoV2 virulence factor^54,55^) – are now recognized to bind the 40S mRNA channel in the same position as LARP1’s RBR-RRM ɑ1. This growing list of factors, along with the coordinated mechanism of LARP1 activation characterized here, reveals the empty mRNA channel of 40S subunits as a privileged site to sense ribosome levels and, more broadly, to regulate ribosome function and cellular homeostasis.

## Methods and Protocols

### Sequence and structure homology analysis

Sequence similarity searches were performed using PSI-BLAST^56^ against the NCBI non-redundant protein database (nr)^57^. Multiple sequence alignments were generated using the MAFFT program^58^ with the local-pair algorithm, combined with the parameters –maxiterate 3000, –op 1.5, and –ep 0.2, and were manually refined based on structural superpositions and profile-profile comparisons. Structure similarity searches were conducted using the DALIlite program^59^ with default parameters and the FOLDSEEK program^27^ tuned for remote homology detection using the following parameters: –c 0.2, –-cov-mode 2, –s 12, –-exhaustive-search 1, –-alignment-type 1. Searches were performed against the PDB database^60^. Phylogenetic trees were constructed using an approximate maximum likelihood model implemented in the FastTree 2.1^61^ program under default parameters. Data processing and visualization were performed using R.

### AlphaFold structural modeling

The AlphaFold2 model of full-length LARP1 (ID: AF-Q6PKG0-F1-v6) was downloaded from the AlphaFold Protein Structure Database^24,62^ (https://alphafold.ebi.ac.uk). For AlphaFold3-based modeling, protein sequences were obtained from UniProt^63^ (https://www.uniprot.org) and imported into the AF3 Server^16^ (https://alphafoldserver.com) either as full-length proteins or with the indicated amino acids deleted. Additional eukaryotic LARP1 homologs with identified RBR-RRM domains were also modeled, using their primary sequences as input. In total, 10 predictions (five models from each of two independent seeds) were generated for each sequence. All models were imported into ChimeraX^64^ version 1.9 (https://www.cgl.ucsf.edu/chimerax/) or Molstar^65^ as.cif files for visualization, analyses, and figure generation. Any manipulations within ChimeraX or Molstar are described in the associated Figure legends.

For predicted aligned error (PAE) plots, matrices were extracted for ten predictions (five models generated from each of two independent seeds) from full_data_*.json files generated by AF3 using the R package jsonlite^66^ and custom scripts written in R. Individual PAE matrices were converted to numeric matrices and stacked into a three-dimensional array representing residue–residue PAE values across predictions. The median PAE value for each residue pair was calculated to generate a consensus matrix. Mean and standard deviation values were also computed for quality control. Median PAE matrices were converted to long format and visualized as heatmaps using ggplot2^67^. Residue indices were plotted with PAE values displayed from 0–30 Å (values greater than 30 Å were masked to improve visualization).

To generate per-residue pLDDT plots, values were extracted for ten predictions (five models generated from each of two independent seeds) from the *.cif file in R using the package bio3d^68^ and custom scripts. For each residue, the mean pLDDT and standard deviation across the ten predications was calculated and plotted across residue indices as a line plot using ggplot2^67^.

To calculate root-mean-square deviation (RMSD), the ChimeraX matchmaker function was used to align residues from each of ten AF3 predictions (five models generated from each of two independent seeds) to residues of the corresponding solved structures (La-wHTH: 400-487, RBR: 660-724, RBR-α1: 673-691, HEAT repeats: 890-999). Using log save, the output was saved as a.txt file and all-atom RMSD values were parsed using custom R scripts.

For TM-scores^69^, the RCSB Pairwise Structure Alignment tool^70^ (https://rcsb.org/alignment) was used to make pairwise comparisons between the RRM domains of ten WT full-length LARP1 AF3 predictions (five models generated from each of two independent seeds; amino acids 640-889) and LARP3 (1ZH5; residues 107-187), LARP4A (6I9B; residues 198-273), LARP6 (2MTG; residues 180-295), and LARP7 (7SLQ; residues 124-380). The structures were aligned using TM-align^71^ and the results were downloaded as.json files using the command line. The results were analyzed using the R package jsonlite^66^ and custom R scripts.

Clashes between the WT LARP1 RBR-RRM/HEAT scaffold and the 40S subunit were determined first by aligning the AF3-predicted α1 helix to the solved structure of the LARP1 RBR bound to the 40S subunit (8XP2; LARP1 residues 673-691). Clashes were determined between ten LARP1 AF3 predictions (five models generated from each of two independent seeds; residues 636-649, 660-724, 797-804, 828-839, and 874-999) and the 40S subunit using the ChimeraX clash function with the parameter overlapCutoff 0.0 and log save to output the resulting clashes as a.txt file. The resulting file was parsed in R using custom scripts. To ensure all atoms were included in the analysis, the AF3 model of the LARP1 RBR-RRM/HEAT scaffold (residues 636-649, 660-724, 797-804, 828-839, and 874-999) was saved as a.pdb and input into R using bio3d^68^ to extract all atoms included in the model.

To calculate the mean percentage of clashed atoms, we counted the atoms of the LARP1 RBR-RRM/HEAT scaffold in each AF3 prediction that had overlap of greater than 0.4 Å (based on the clashscore metric^72^) with an atom in the 40S subunit. The percentage was calculated out of the total number of atoms in the LARP1 RBR-RRM/HEAT scaffold. The mean percentage and standard deviation were calculated and visualized using ggplot2^67^.

To calculate the mean overlap per atom in the LARP1 RBR-RRM/HEAT scaffold, we first defined the full set of possible interactions between LARP1 and the 40S subunit as the union of every LARP1 atom-40S atom pair in at least one of the ten models and all LARP1 RBR-RRM/HEAT atoms. For each AF3 prediction, the full set of possible interactions was assessed: LARP1 atoms that did not participate in any interaction and interactions that were not present for that AF3 prediction (but were present in the full set) were retained with an overlap set to zero. The mean overlap for each interaction of the full set across each of the ten AF3 models was calculated. For LARP1 atoms with multiple interactions with atoms in the 40S subunit, the interaction with the greatest mean overlap was retained for plotting. The mean overlap was plotted for each atom in the LARP1 RBR-RRM/HEAT scaffold using ggplot2^67^.

### Cell culture

HEK293T WT (ATCC CRL-3216) and LARP1-KO^12^ cell lines were cultured in DMEM (ThermoFisher 11995073) supplemented with 10% FBS (ThermoFisher A5670701). All experiments were performed in DMEM with 10% FBS unless otherwise specified and begun 24 hours after seeding to allow time for cells to attach. Cells were passaged with 0.25% Trypsin-EDTA (ThermoFisher 25200114), maintained at 37°C and 5% CO_2_, and regularly tested negative for mycoplasma.

### LARP1 mutant plasmid construction

The pLJC1-LARP1 plasmid expressing the LARP1 1019 amino acid isoform (ENSEMBL LARP1-201) behind a CMV promoter^12^ was linearized by PCR and a gene block (IDT) introducing the first 77 amino acids of the LARP1 1096 amino acid isoform (ENSEMBL LARP1-204) was integrated by Gibson Assembly (NEB E2611L) according to the manufacturer’s instructions to generate the WT-LARP1 plasmid construct. This plasmid was used as the backbone for all LARP1 full-length mutant constructs. The *β*1-P, *β*1-del, and *β*3-del plasmids, were constructed using site-directed mutagenesis with primers introducing the appropriate amino acid substitutions or deletions. The RBRmut-1, RYW-mut, RBRmut-2, HEAT-mut, Phos-null, Phos-mim, and GFP plasmids, as well as double mutant constructs, were constructed by Gibson Assembly, integrating a gene block or PCR-amplified product with the appropriate amino acid substitutions into the appropriate linearized backbone.

N-terminally truncated LARP1^574-1096^ plasmids were constructed as described above but using the WT-LARP1^574-1096^ plasmid ^12^ as the backbone.

### Luciferase reporter assay

TOP and non-TOP reporter constructs containing the RPL39 or UBL5 promoter and 5’UTR, respectively, followed by an nLuc-PEST sequence were obtained from^14^. LARP1-KO cells were seeded in 96-well plates (Corning 3610) at 1 x 10^4^ cells per well. After 24 hours, cells were transfected with 10 ng of the appropriate LARP1 or GFP plasmid and 25 ng of the appropriate reporter plasmid using Lipofectamine 3000 (ThermoFisher L3000015) according to the manufacturer’s instructions. The following day, cells were treated with 300 nM Torin1 (Selleck Chem S2827) or an equivalent volume of DMSO (1%) for 1 hour, then subjected to the NanoGlo Dual-Luciferase Reporter Assay (Promega N1620) according to the manufacturer’s instructions. Luciferase luminescence was recorded with the Synergy H1 Microplate Reader (BioTek). Nano luciferase (Nluc) was normalized to Firefly luciferase (FFluc) for each sample. All analysis was performed using custom R scripts and visualized using ggplot2^67^.

### Immunoblotting

Cells were seeded in 6-well plates (ThermoFisher 140675) at 2 x 10^5^ cells per well. After 24 hours, cells were transfected with the appropriate LARP1 or GFP plasmid (150 ng full-length LARP1 or GFP plasmid or 200 ng N-terminally truncated LARP1^574-1096^ or GFP plasmid) using Lipofectamine 3000 (ThermoFisher L3000015) according to the manufacturer’s instructions.

The following day, the media was refreshed and 2 hours later, cells were washed once with 1 mL PBS (Gibco 10010023), then collected with 100 µL 0.25% Trypsin-EDTA (ThermoFisher 25200114) and 1400 µL media. Cells were pelleted by centrifugation at 350 x g for 3 mins, then lysed in 110 µL protein lysis buffer (50 mM HEPES pH 7.4, 100 mM KOAc, 15 mM Mg(OAc)_2_, 5% glycerol, 0.25% Triton X-100 (MilliporeSigma T2984), 360 µM emetine dihydrochloride (MilliporeSigma E2375), 1 mM TCEP (Gold Biotechnology TCEP2), 8 U/mL TURBO DNase (ThermoFisher AM2239), 1x Halt protease and phosphatase inhibitor cocktail (ThermoFisher 78445). Cell lysates were clarified by centrifugation at 8,000 x g for 5 mins at 4°C, then normalized to total protein content by BCA assay (ThermoFisher 23225). Cell lysates were boiled at 95°C for 5 mins in 1x Laemmeli buffer prior to loading on the gel.

Gel electrophoresis was performed with 4-10 ug of sample using 4-20% Criterion TGX gels (BioRad 5671095) and transferred to a PVDF membrane (BioRad 12023927) using a 30 min standard transfer on the Trans-Blot Turbo Transfer System (BioRad). Membranes were blocked with 5% milk (Santa Cruz Biotechnology sc-2324) in TBST (Tris-buffered saline with 1% Tween 20) for 1 hour, then incubated in primary antibody overnight at the appropriate dilution in 5% milk in TBST at 4°C. The following day, membranes were washed 3x with TBST, incubated in the appropriate secondary antibody at 1:5000 dilution in 5% milk in TBST for 1 hour, then washed 3x with TBST again. Membranes were incubated in chemiluminescent reagent (ThermoFisher 34580X4; ThermoFisher 34096X4) then imaged using a ChemiDoc imaging system (BioRad). For FLAG-LARP1 immunoblotting, secondary antibody incubation was omitted. Membranes were probed with the following antibodies at the following dilutions: RACK1 (1:1000, Cell Signaling 5432S); RPS3 (1:1000, Abcam ab140688); RPS24 (1:1000, Abcam ab196652); 4E-BP1 (1:1000, Cell Signaling 9644S); Vinculin (1:2000, Santa Cruz Biotechnology sc-73614); Actin (1:1000, Cell Signaling 4967S); LARP1 (1:1000, Cell Signaling 70180S; Supplemental Figures S8C, 9D, S13A); LARP1 (1:1000, Abcam ab86359; Supplemental Figures S8A, S9B); FLAG-HRP (1:10000, MilliporeSigma A8592); Rabbit IgG-HRP (Cell Signaling 7074S); Mouse IgG-HRP (Cell Signaling 7076S)

### Ribosome binding co-immunoprecipitation

LARP1-KO cells were seeded at 1 x 10^6^ cells per dish in 10 cm dishes (Corning 430167). After 24 hours, 750 ng of the appropriate N-terminally truncated LARP1^574-1096^ or GFP plasmid was transfected using Lipofectamine 3000 (ThermoFisher L3000015) according to the manufacturer’s instructions. After approximately 16 hours, the media was refreshed and 1 hour later, cells were treated with 300 nM Torin1 (Selleck Chem S2827) or an equivalent volume of DMSO (0.03%) for 1 hour. At the time of lysis, cells were washed with 4 mL PBS (Gibco 10010023) and 250 µL immunoprecipitation (IP) lysis buffer was added directly to the cells (<u>IP</u> <u>lysis buffer</u>: 50 mM HEPES pH 7.4, 100 mM KCl (Invitrogen AM9640G), 15 mM MgCl_2_ (Invitrogen AM9530G), 5% glycerol, 0.25% Triton X-100 (MilliporeSigma T2984), 360 µM emetine dihydrochloride (MilliporeSigma E2375), 20 U/mL SUPERaseIN (ThermoFisher AM2694), 8 U/mL TURBO DNase (ThermoFisher AM2239), 1x Halt protease and phosphatase inhibitor cocktail (ThermoFisher 78445)). Cell lysates were collected in IP lysis buffer by scraping cells directly from the dish, pipetting to homogenize, and centrifugation at 8000 x g for 5 min at 4°C. Clarified lysates were normalized to protein content by BCA assay (ThermoFisher 23225) and 5% (v/v) was set aside and diluted 1:20 in IP lysis buffer as input. The remaining lysates were added to 30 µL of anti-FLAG M2 magnetic beads (MilliporeSigma M8823) washed twice in IP lysis buffer. Lysates were incubated with beads for 1 hour at 4°C on a nutator to allow binding to beads. Beads were washed 3x in wash buffer 1 and 3x in wash buffer 2 for 5 mins at 4°C on a nutator and collected each time using a magnetic tube rack (<u>wash buffer 1</u>: 50 mM HEPES pH 7.4, 100 mM KCl, 15 mM MgCl_2_, 5% glycerol, 0.1% Triton X-100, 1x Halt protease and phosphatase inhibitor cocktail; <u>wash buffer 2</u>: 50 mM HEPES pH 7.4, 100 mM KCl, 15 mM MgCl_2_, 0.05% Triton X-100, 1x Halt protease and phosphatase inhibitor cocktail). Following all washes, beads were resuspended in 60 µL elution buffer (wash buffer 2 with 400 µg/mL FLAG peptide (MilliporeSigma F3290)) and incubated for at least 1 hour at 4°C on a nutator. Input and elution samples were boiled at 95°C for 5 mins in 1x Laemmeli buffer prior to loading on the gel. Samples (0.05% input; 15% elution) were loaded onto 4-20% Criterion TGX gels (BioRad 5671095) and processed as described in “Immunoblotting” with the following antibodies: RACK1, RPS3, RPS24, 4E-BP1, Vinculin, and FLAG.

### 40S subunit purifications

HEK293T WT cells were seeded at 5 x 10^6^ cells per dish in five 15 cm plates (FisherScientific FB012925). After 24 hours, cells were washed with 8 mL PBS (Gibco 10010023) and 2 mL of ribosome lysis buffer was added directly to the plate (<u>ribosome lysis buffer</u>: 50 mM HEPES pH 7.4, 100 mM KCl (Invitrogen AM9640G), 5 mM MgCl_2_ (Invitrogen AM9530G), 5% glycerol, 0.5% Triton X-100 (MilliporeSigma T2984), 1 mM dithiothreitol (DTT; Gold Biotechnology DTT), 20 U/mL SUPERaseIN (ThermoFisher AM2694), 8 U/mL TURBO DNase (ThermoFisher AM2239), 1x Halt protease and phosphatase inhibitor cocktail (ThermoFisher 78445)). Cell lysates were collected in ribosome lysis buffer by scraping directly from the dish, pipetting briefly to homogenize, then centrifugation at 8000 x g for 5 min at 4°C.

To isolate ribosomes, 1.5 mL cell lysate was layered over a 1 mL sucrose cushion in 3.5 mL polycarbonate ultracentrifuge tubes (Beckman Coulter 349622) and ultracentrifuged at 100,000 rpm for 1 hour at 4°C using a TLA100.3 rotor (Beckman Coulter 349481) (<u>sucrose</u> <u>cushion</u>: 50 mM HEPES pH 7.4, 500 mM KCl, 25mM MgCl_2_, 1 mM DTT, 0.5 M sucrose). Following ultracentrifugation, ribosomes were resuspended in 100 uL of subunit separation buffer then incubated with 2 mM puromycin (MilliporeSigma P7255) for 15 mins on ice followed by 10 mins at 37°C with gentle shaking (<u>subunit separation buffer</u>: 50 mM HEPES pH 7.4, 500 mM KCl, 2 mM MgCl_2_, 2 mM DTT). Samples were layered over 10-35% sucrose gradients (150 µL per gradient) constructed as follows: 6 mL of 10% (w/v) sucrose buffer was pipetted into SW41 polypropylene ultracentrifuge tubes (Seton Scientific 7030), then 6 mL of 35% (w/v) sucrose buffer was added to the bottom of the tube using a syringe and cannula, and mixed on a Biocomp Gradient Master (<u>sucrose buffer</u>: 50 mM HEPES pH 7.4, 500 mM KCl, 2 mM MgCl_2_, 2 mM DTT, appropriate concentration sucrose). Gradients were ultracentrifuged at 40,000 rpm for 1:45 hours at 4°C using a SW41 rotor (Beckman Coulter 331362). Ultracentrifuged gradients were fractionated and A260 absorbance was measured on a Biocomp Piston Gradient Fractionator according to the manufacturer’s instructions. Fractions containing 40S subunits were pooled and pelleted in 3.5 mL polycarbonate ultracentrifuge tubes by ultracentrifugation at 100,000 rpm for 2 hours at 4°C using a TLA100.3 rotor. The supernatant was discarded and 40S subunits were resuspended in storage buffer (20 mM HEPES pH 7.4, 50 mM NaCl, 50 mM KCl, 5 mM MgCl_2_, 2 mM DTT). A260 measurement was taken using a Nanodrop (ThermoFisher) to determine the final concentrations and 40S subunits were stored at –80°C.

### LARP1 purifications

WT, RBRmut-2, *β*1-P, and HEAT-mut LARP1^574-1096^ was purified from adherent LARP1-KO HEK293T cells using a similar protocol to ^12^ with slight modifications. Cells were seeded to 6.5 x 10^6^ cells per plate in five 15 cm plates (FisherScientific FB012925). The next day, cells were transfected with 25 µg per plate of the appropriate N-terminally truncated LARP1^574-1096^ plasmid using Lipofectamine 3000 (ThermoFisher L3000015) according to the manufacturer’s instructions and allowed to grow for two days with media refreshed on day one post-transfection. On day two post-transfection, cells were scraped from the plate and pelleted in 2 mL cold lysis buffer per plate (<u>lysis buffer</u>: 50 mM HEPES pH 7.4, 100 mM NaCl, 1% Triton X-100 (MilliporeSigma T2984), 2 mM EDTA, 5% glycerol, 1x Halt protease and phosphatase inhibitor cocktail (ThermoFisher 78445), 8 U/mL TURBO DNase (ThermoFisher AM2239), 10 U/mL benzonase (MilliporeSigma E1014)). Lysates were clarified by centrifugation at 16,000 x g for 5 min at 4°C. Clarified lysates were incubated with 10 µL of packed anti-FLAG M2 affinity beads (MilliporeSigma A2220) per plate of cells for 2 hours at 4°C with gentle rotation. Following incubation, beads were transferred to a chromatography column on ice and washed twice with 1 mL cold lysis buffer, twice with 5 mL cold wash buffer 1 (50 mM HEPES pH 7.4, 1 M NaCl, 0.2% Triton X-100, 2 mM EDTA, 5% glycerol), and twice with 5 mL cold wash buffer 2 (50 mM HEPES pH 7.4, 1 M NaCl, 2 mM EDTA, 5% glycerol). Beads were eluted in 500 µL of cold elution buffer containing 500 µg/mL FLAG peptide (MilliporeSigma F3290) (<u>elution buffer</u>: 50 mM HEPES pH 7.4, 1 M NaCl, 5% glycerol, 500 µg/mL FLAG peptide) for 30 min at 4°C with gentle rotation. Purified, eluted protein was concentrated twice to 100 uL in storage buffer using a 30K MWCO PES filter (ThermoFisher 88502) to remove FLAG peptide and replace NaCl with KCl (<u>storage</u> <u>buffer</u>: 50 mM HEPES pH 7.4, 200 mM KCl, 10% glycerol). Concentrated protein was flash frozen and stored at –80°C.

### Oligonucleotide capping and labeling

Purified TOP (ppp-CUCUUCCGCCGU) and non-TOP (ppp-GAGAAGGGCCGU) RNA oligonucleotides containing a 5’ triphosphate modification were custom ordered from GeneWiz. For *in vitro* TOP binding assays (Figures 3A-E and Supplemental Figures S12D-G), the oligonucleotides were capped and labeled with GTP[ɑ-^32^P] (Revvity BLU006H250UC) using the Vaccinia Capping System (NEB M2080S), with modifications to the protocol to increase yield. Five hundred pmol of RNA was incubated with 1x capping buffer, 100 µM S-adenosylmethionine, 15 µL of GTP[ɑ-^32^P], and 10 µL of vaccinia capping enzyme for a 50 µL total reaction for 1 hour at 37°C. Capped, radiolabeled RNA was precipitated by 2-propanol precipitation, washed in 80% ethanol, and resuspended in RNase-free water. RNA was run over a G-25 column (MilliporeSigma GE27-5325-01) to remove free GTP[ɑ-^32^P]. Capping efficiency was assessed by TBE-urea polyacrylamide gel stained with SYBR gold (ThermoFisher S11494).

For *in vitro* 40S binding assays (Figure 3F), the oligonucleotides were capped with unlabeled GTP using the Vaccinia Capping System according to the manufacturers instructions. Capped RNA was precipitated by 2-propanol precipitation, washed in 80% ethanol, resuspended in RNase-free water, and run over a G-25 column to remove free GTP. Capping efficiency was assessed by TBE-urea polyacrylamide gel stained with SYBR gold.

### *In vitro* binding assays

For *in vitro* TOP binding assays (Figures 3A-E and Supplemental Figures S12D-G), 0.2 µL of packed anti-FLAG M2 magnetic beads (MilliporeSigma M8823) per sample was equilibrated in binding buffer (<u>binding buffer</u>: 50 mM HEPES pH 7.4, 200 mM KCl (Invitrogen AM9640G), 2.5 mM MgCl_2_ (Invitrogen AM9530G), 5% glycerol, 0.5 mg/mL BSA (NEB B9200S), 0.01% Triton X-100 (MilliporeSigma T2984)). Five nM of purified LARP1^574-1096^ was bound to beads for 5 minutes, followed by addition of 5 nM capped, radiolabeled TOP or non-TOP oligo, and varying amounts of purified 40S subunits. Samples were incubated overnight at 4°C with gentle rotation. The next day, samples were washed once in binding buffer and eluted in 10 µL of binding buffer supplemented with 500 µg/mL FLAG peptide (MilliporeSigma F3290) for 1 hour at 4°C with gentle rotation. Following elution, 5 µL of RNA loading buffer (NEB B0363S) was added to each sample and samples were boiled for 1 min at 95°C. Samples were run on a 15% TBE-urea polyacrylamide gel at 250 V for 1 hour. The gel was vacuum-dried and exposed to a phosphor screen overnight. Following exposure, the phosphor screen was imaged on an Amersham Typhoon phosphoimager (Cytiva). Autoradiograph band intensity was quantified using ImageJ ^73^. Background was subtracted from raw images using a 500 pixel rolling ball radius and band intensity was quantified using the Gel Analyzer tools, then output as a.csv. Plots were generated in R using custom scripts.

For *in vitro* 40S binding assays (Figure 3F), 1 µL of packed anti-FLAG M2 magnetic beads per sample was equilibrated in binding buffer. Fifty nM of purified LARP1^574-1096^ was bound to beads for 5 minutes, followed by addition of 50 nM of purified 40S subunits and varying amounts of capped, unlabeled TOP or non-TOP oligo. Samples were incubated for 1 hour at 4°C with gentle rotation. Following incubation, samples were washed once in binding buffer and eluted by boiling at 95°C for 5 min in 1x Laemmli buffer. Samples were loaded onto 4-20% Criterion TGX gels (BioRad 5671095) and processed as described in “Immunoblotting” with antibodies probing for 40S subunit ribosomal proteins (RACK1 and RPS24).

### Sucrose gradient analysis

LARP1-KO cells were seeded in 10 cm dishes (Corning 430167) at 1 x 10^6^ cells per dish. After 24 hours, 750 ng of the appropriate LARP1 or GFP plasmid was transfected with Lipofectamine 3000 (ThermoFisher L3000015) according to the manufacturer’s instructions. After approximately 16 hours, the media was refreshed and 1 hour later, cells were treated with DMSO (0.03%) for 1 hour. At the time of lysis, cells were washed with 4 mL of ice-cold PBS (Gibco 10010023) supplemented with 360 µM emetine dihydrochloride (MilliporeSigma E2375) and 250 µL gradient lysis buffer was added directly to the cells (<u>gradient lysis buffer</u>: 50 mM HEPES pH 7.4, 100 mM KOAc, 15 mM Mg(OAc)_2_, 5% glycerol, 0.25% Triton X-100 (MilliporeSigma T2984), 360 µM emetine dihydrochloride, 1 mM TCEP (Gold Biotechnology TCEP2), 20 U/mL SUPERaseIN (ThermoFisher AM2694), 8 U/mL TURBO DNase (ThermoFisher AM2239), 1x Halt protease and phosphatase inhibitor cocktail (ThermoFisher 78445)). Cells were collected by scraping directly from the dish, pipetting gently to homogenize, then transferred to tubes on ice. Cell lysates were clarified by centrifugation at 8,000 x g for 5 mins, then normalized to total RNA content by QUBIT Broad Range assay (Invitrogen Q10211).

Sucrose gradients were prepared as follows: 6 mL of 10% (w/v) sucrose buffer was pipetted into SW41 polypropylene ultracentrifuge tubes (Seton Scientific 7030), then 6 mL of 50% (w/v) sucrose buffer was added to the bottom of the tube using a syringe and cannula, and mixed on a Biocomp Gradient Master (<u>sucrose buffer</u>: 50 mM HEPES pH 7.4, 100 mM KOAc, 5 mM Mg(OAc)_2_, 1 mM TCEP, 180 µM emetine dihydrochloride, 4 U/mL SUPERaseIN, appropriate concentration sucrose). Gradients were ultracentrifuged at 40,000 rpm for 1:30 hours at 4°C using a SW41 rotor (Beckman Coulter 331362). Ultracentrifuged gradients were fractionated and A260 absorbance was measured on a Biocomp Piston Gradient Fractionator according to the manufacturer’s instructions. Sucrose gradient fractions (250 µL) were transferred to 4x volumes of TRIzol reagent (Invitrogen 15596026) supplemented with 100 pg of *β*-galactosidase mRNA spike-in per fraction (TriLink Biotechnologies L-7608-100).

### Steady state qPCR

LARP1-KO cells were seeded in 6-well plates (ThermoFisher 140675) at 2×10^5^ cells per well. After 24 hours, cells were transfected with 150 µg of the appropriate LARP1 or GFP plasmid using Lipofectamine 3000 (ThermoFisher L3000015) according to the manufacturer’s instructions. The following day, the media was changed to media treated with DMSO (0.03%) for 24 hours. At the time of lysis, cells were washed once with 1 mL PBS (Gibco 10010023), then collected with 100 µL 0.25% Trypsin-EDTA (ThermoFisher 25200114) and 1400 µL media. Cells were pelleted by centrifugation at 350 x g for 3 mins, then the pellet was lysed in 110 µL gradient lysis buffer (<u>gradient lysis buffer</u>: 50 mM HEPES pH 7.4, 100 mM KOAc, 15 mM Mg(OAc)_2_, 5% glycerol, 0.25% Triton X-100 (MilliporeSigma T2984), 360 µM emetine dihydrochloride (MilliporeSigma E2375), 1 mM TCEP (Gold Biotechnology TCEP2), 20 U/mL SUPERaseIN (ThermoFisher AM2694), 8 U/mL TURBO DNase (ThermoFisher AM2239), 1x Halt protease and phosphatase inhibitor cocktail (ThermoFisher 78445)). Cell lysates were clarified by centrifugation at 8,000 x g for 5 mins at 4C, then immediately transferred to 1 mL TRIzol reagent (Invitrogen 15596026) and flash frozen.

### RNA extractions and RT-qPCR

RNA was extracted from samples (sucrose gradient fractions or whole cell lysates) stored in TRIzol reagent (Invitrogen 15596026) according to the manufacturer’s instructions. At the final step, isolated RNA was resuspended in 25 µL nuclease-free water. Prior to reverse transcription (RT), 8 µL RNA from sucrose gradient fraction pools or 2 µg RNA from whole cell lysate was heated at 70°C for 5 mins, then immediately placed on ice. Reverse transcription was performed in 40 µL reactions with the iScript cDNA synthesis kit (BioRad 1708891) according to the manufacturer’s instructions. Following RT, cDNA was diluted by adding 170 µL nuclease-free water. At this step, diluted cDNA from sucrose gradient fractions was pooled as follows: 40 µL cDNA from fractions 1-3 was pooled with 80 µL nuclease-free water, 40 µL cDNA from fractions 1-5 was pooled. qPCR was performed by assembling reactions containing 3.5 µL diluted cDNA with 1.5 µL of 3 µM forward and reverse primer mix targeting the gene of interest (450nM final primer concentration) and 5 µL iTaq Universal SYBR Green Supermix (BioRad 1725125) using a QuantStudio 6 RT-qPCR system with the following protocol: 95°C for 1 min, 40 x (95°C for 10 secs, 60°C for 30 secs) then 95°C for 15 secs, 60°C for 1 min, then to 95°C at 0.05°C/s for 15 secs. All RT-qPCR reactions were performed in technical triplicate. Samples were probed with the following primers. All primers were verified as yielding a single amplified band by gel electrophoresis.

BGAL_F: AGGACATGTGGCGGATGAG

BGAL_R: GGCTGAAGTCGTCGTTGAAC

RACK1_F: GCTGATGGCCAGACTCTGTT

RACK1_R: TTCTAGCGTGTGCCAATGGT

RPS3A_F: CCCGAGAGGTGCAGACAAAT

RPS3A_R: GCTCCATGAGCTTTCCCAATTC

RPS8_F: GGTACGAGTCCCACTATGCG

RPS8_R: GAAGCGATGCACGCAAGAAG

ACTB_F: ACGTTGCTATCCAGGCTGTG

ACTB_R: GAGGGCATACCCCTCGTAGA

RPL5_F: CAGCGTATGCACACGAACTG

RPL5_R: ACCTATTGAGAAGCCTGCGG

RPL23_F: GATGTCGAAGCGAGGACGTG

RPL23_R: CTGTTCAGCCGTCCCTTGAT

MYC_F: AGTGGAAAACCAGCAGCCTC

MYC_R: TTCTCCTCCTCGTCGCAGTA

For each experiment, a standard curve of cDNA diluted two-fold seven times was probed with each primer set and was used to interpolate the relative cDNA quantity for each sample. For sucrose gradient fraction pools, qPCR reactions targeting the *β*-galactosidase mRNA spike in (BGAL) were performed to control for differences arising from RNA extraction or RT efficiency. The relative quantity of the gene of interest for each cDNA sample was normalized to that of BGAL. The proportion of the gene of interest in the polysome pool was calculated relative to the total in both pools. All analysis was performed using custom R scripts and visualized using ggplot2.

### Cell growth quantification

LARP1-KO cells were seeded in 12-well plates (Corning 3513) at 7.5 x 10^5^ cells per well. After 24 hours, 75 ng of the appropriate LARP1 or GFP plasmid was transfected using Lipofectamine 3000 (ThermoFisher L3000015) according to the manufacturer’s instructions. After 16 hours, the media was changed to the appropriate conditions. For growth assays (Figure 4C-D), the media was refreshed (and supplemented with DMSO at 0.03%) and imaging began in the CellCyte X (Cytena) 2-4 hours later. For recovery assays (Supplemental Figure S13E), the media was changed to DMEM (ThermoFisher 11995073) without FBS for 24 hours, then FBS (ThermoFisher A5670701) was added to 10% and imaging began in the CellCyte X 2-4 hours later. For each of three biological replicates, each LARP1 mutant sample was plated in triplicate (3 wells) with 4 images (Enhanced contour) taken per well with classifier-based autofocus enabled. Images were taken every 6 hours for 36-48 hours.

After imaging was complete, analysis was conducted on the CellCyte X software using the following parameters recipe: cell confluence; contrast sensitivity 50 a.u.; smoothing 2 a.u.; filled hole size 100 µm^2^; minimum object size 100 µm^2^. Raw percent confluence values for each image were input to a custom R script and used to calculate the mean percent confluence across each condition for each time point, then plotted as a growth curve. To calculate doubling time, the mean percent confluence for each well across the time course was fit to a logistic curve 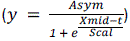 using the SSlogis function in R. The maximum of the derivative of the fit was calculated as the maximum growth rate (in percent confluence/hour; max_rate), from which the doubling time was calculated using the equation 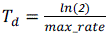. The mean doubling time across wells for each LARP1 mutant was averaged for each biological replicate. All analysis was performed using custom R scripts and visualized using ggplot2.

## Statistical analysis

All statistical analyses were performed in R. Standard deviations were calculated using the stats package and standard error of the mean values were calculated from the standard deviation. ANOVA tests (one-way and two-way) were performed using the stats package followed by a correction for multiple comparisons: Dunnett’s correction for multiple comparisons was applied using the package emmeans with parameters method = “trt.v.ctrl” and adjust = “mvt”) and using the WT sample as the reference; Tukey’s correction for multiple comparisons was applied using the package emmeans and the relevant comparisons are shown. For two-way ANOVA tests, Dunnett’s correction was applied within each treatment group. P values were extracted and reported as follows: n.s. not significant, * p ≤ 0.05, ** p ≤ 0.01, *** p ≤ 0.001, **** p ≤ 0.0001.

## Data Availability

All data are available in the manuscript or supplemental materials. ‘This study includes no data deposited in external repositories. No code is necessary for interpretation or replication of the study results. All materials used in this study are freely available upon request to the corresponding author.

## Author contributions

J.A.S., P.E.W., and R.G. conceptualized the project. P.E.W. and J.A.S. performed experiments. A.M.B. performed phylogenetic analyses. J.A.S. wrote the manuscript with input from all study authors. J.A.S., P.E.W., A.M.B, L.A., and R.G. reviewed and edited the manuscript. R.G. oversaw and funded the project.

## Acknowledgments

We thank Carson Thoreen for providing LARP1-KO cell lines and pLJC1-LARP1 and truncated LARP1^497-1019^ (here called LARP1^574-1096^) plasmids. We thank members of the Green and Aravind labs for helpful comments throughout the course of this study. This work utilized the computational resources of the NIH HPC Biowulf cluster (https://hpc.nih.gov).

## Funding

J.A.S. is supported by the National Institutes of Health MSTP grant (T32GM136577). P.E.W. is supported by the National Institutes of Health training grant (T32GM144272). A.M.B. and L.A. are supported by the Intramural Research Program of the National Institutes of Health (NIH). R.G. is supported by the Howard Hughes Medical Institute.

**Supplemental Figure S1.**
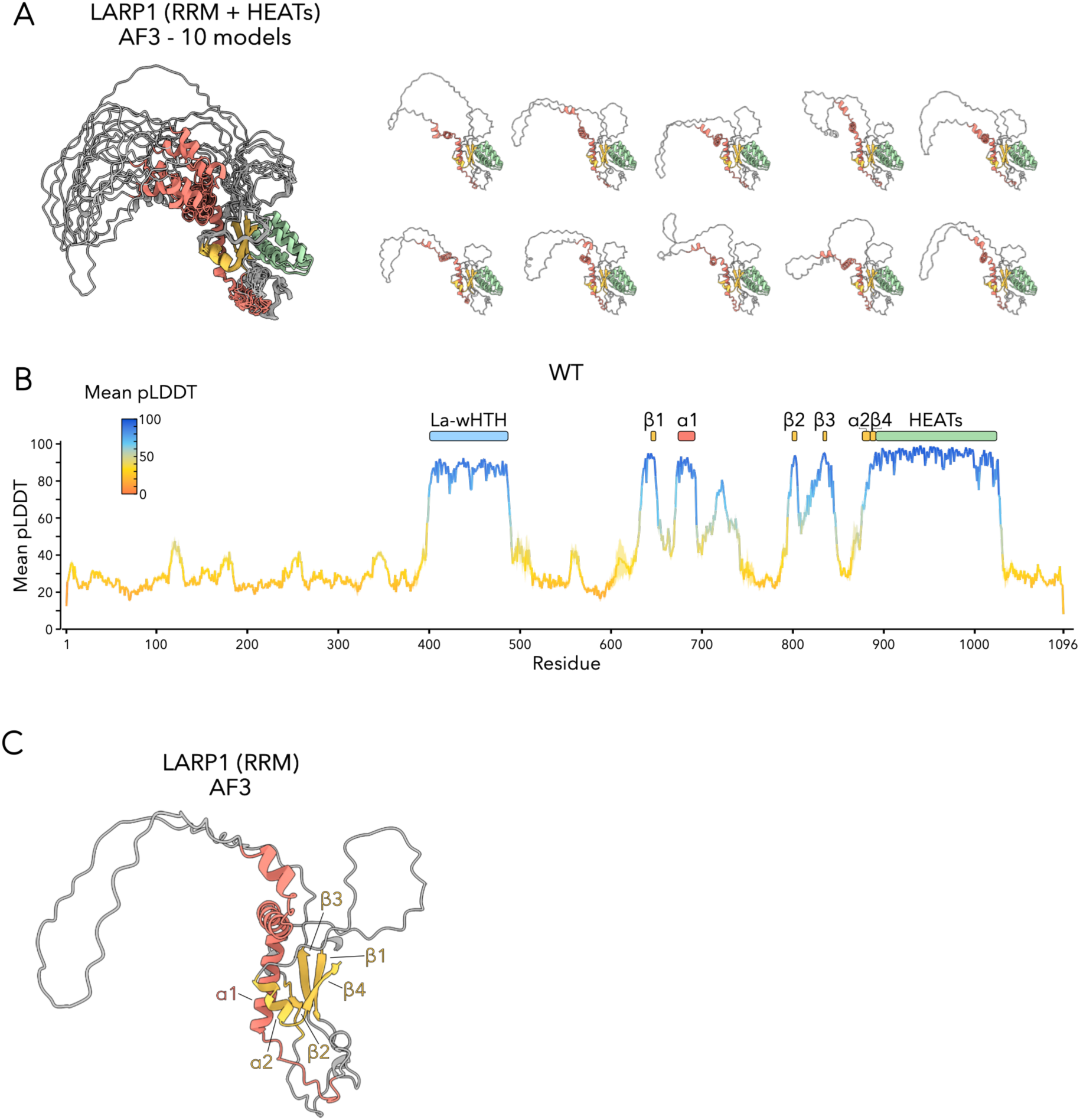
Characterization of the LARP1 AlphaFold3 model. Related to Figure 1. **A)** Ten AF3 models (five models from each of two independent seeds) of full-length LARP1 overlaid (left) and split into individual models (right). Models are aligned to the HEAT repeats (890-999). The region spanning the RBR-RRM to HEAT repeats (640-999) is shown for clarity. **B)** Mean per-residue pLDDT plot for WT-LARP1 from ten AF3 predictions (five models generated from each of two independent seeds). The line plot represents the mean pLDDT scores and the shaded ribbon represents ± one standard deviation. High pLDDT scores (blue) indicate high confidence in the local folding of the La-wHTH, RBR-RRM, and HEAT repeat domains. **C)** AF3 model of the LARP1 RBR-RRM with predicted structured elements labeled.

**Supplemental Figure S2.**
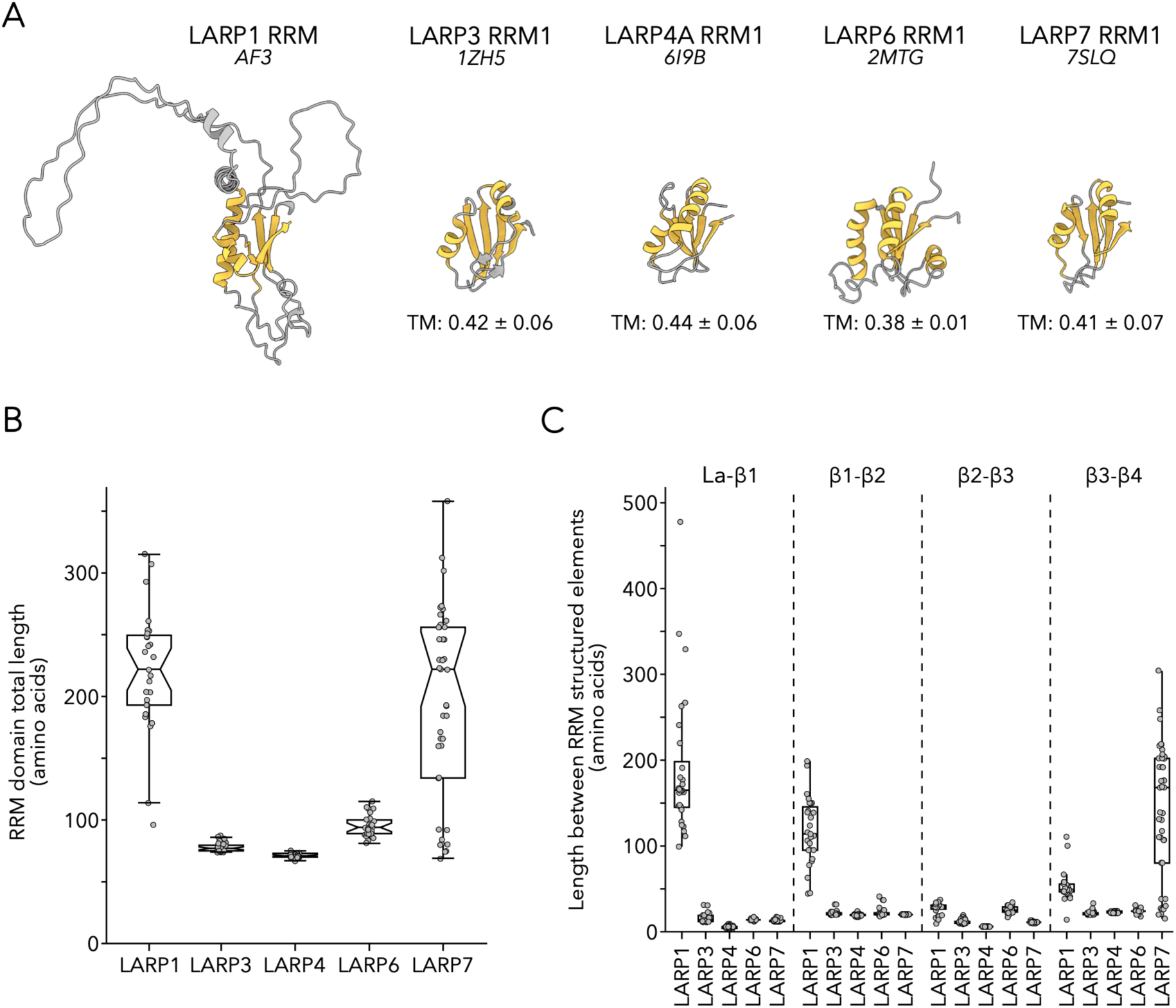
Characterization of the LARP1 RBR-RRM. Related to Figure 1. **A)** Representative AF3 model of LARP1 RBR-RRM, compared to solved structures of RRMs from LARP3 (1ZH5), LARP4 (6I9B), LARP6 (2MTG), and LARP7 (7SLQ). The mean TM-score^69^ ± one SD of ten AF3 models (five models from each of two independent seeds) of the LARP1 RBR-RRM compared to the respective RBR-RRM of each LARP family protein is reported below the depicted structure. **B)** Boxplots depicting total amino acid length of the RBR-RRM domain immediately C-terminal to the La-wHTH domain across LARP families, as measured from the first residue of the RRM *β*1 strand to the final residue of *β*4. **C)** Boxplots depicting amino acid lengths of regions connecting designated secondary structure elements in the core RBR-RRM domain. The first set of boxplots depicts the distances between the end of the La-wHTH (La) domain and the start of the RRM *β*1 strand.

**Supplemental Figure S3.**
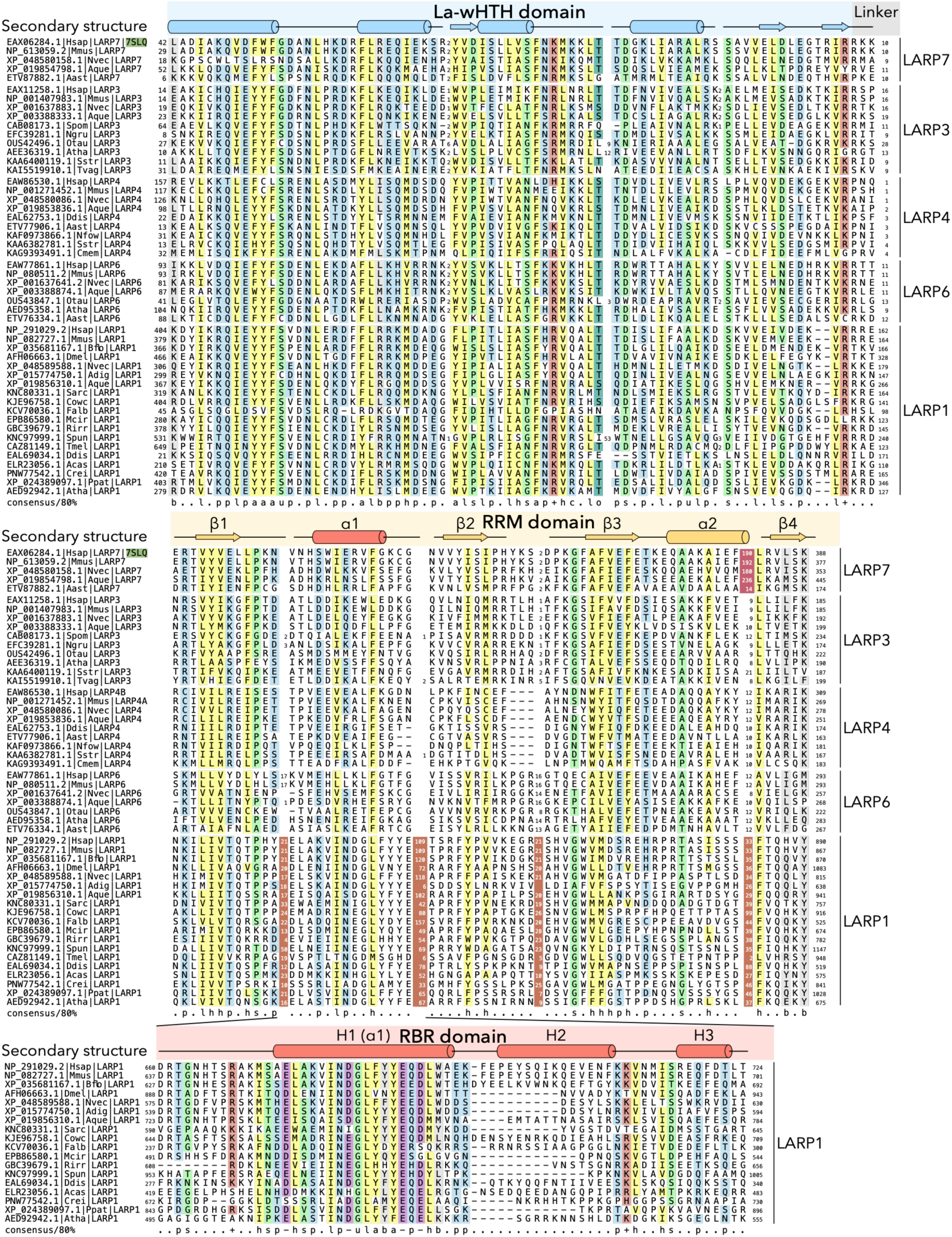
Multiple sequence alignment of the La-wHTH and C-terminal RRM domains across LARP families. Condensed version of the multiple sequence alignment underlying the phylogenetic tree in Figure 1B. The La-wHTH domain is included as an anchor, to emphasize the relative positioning of the RRM domains across families. LARP families are labeled to the right of the alignment. Individual sequences are labeled to the left with GenBank accession number, organism abbreviation, and LARP family name, separated by vertical bars. PDB identifiers, shaded in green, are included where appropriate. Secondary structure elements at top of the alignment are represented as follows: disordered regions as black lines, *β*-strands as extended arrows, and α-helices as cylinders. Numbers to the left and right of the alignment correspond to amino acid position. Insert regions are replaced with numbers indicating the amino acid length of the insert. Inserts within the RBR-RRM domain are shaded in red. Inset at the bottom provides alignment of the RBR of LARP1. Residues are colored according to amino acid conservation, as indicated by the consensus line below the alignment, abbreviations are as follows: h, hydrophobic (yellow); l, aliphatic (yellow); a, aromatic (yellow); s, small (green); u, tiny (green); –, negatively charged (purple); p, polar (blue); c, charged (blue); +, positively charged (red); b, big (gray). Organism abbreviations as follows: Hsap: *Homo sapiens*; Acas*: Acanthamoeba castellanii*; Ddis: *Dictyostelium discoideum*; Ngru: *Naegleria gruberi*; Sstr*: Streblomastix strix*; Spom: *Schizosaccharomyces pombe*; Dmel: *Drosophila melanogaster*; Bflo: *Branchiostoma floridae*; Mmus: *Mus musculus*; Nvec: *Nematostella vectensis*; Aque: *Amphimedon queenslandica*; Falb: *Fonticula alba*; Crei: *Chlamydomonas reinhardtii*; Atha: *Arabidopsis thaliana*; Otau: *Ostreococcus tauri*; Tvag: *Trichomonas vaginalis*; Aast: *Aphanomyces astaci*; Cmem: *Carpediemonas membranifera*; Mcir: *Mucor circinelloides*; Nfow: *Naegleria fowleri*; Akon: *Acrasis kona*; Ppat: *Physcomitrium patens*; Adig: *Acropora digitifera*; Rirr*: Rhizophagus irregularis*; Sarc: *Sphaeroforma arctica*; Cowc: *Capsaspora owczarzaki*; Spun: *Spizellomyces punctatus*; Tmel: *Tuber melanosporum*.

**Supplemental Figure S4.**
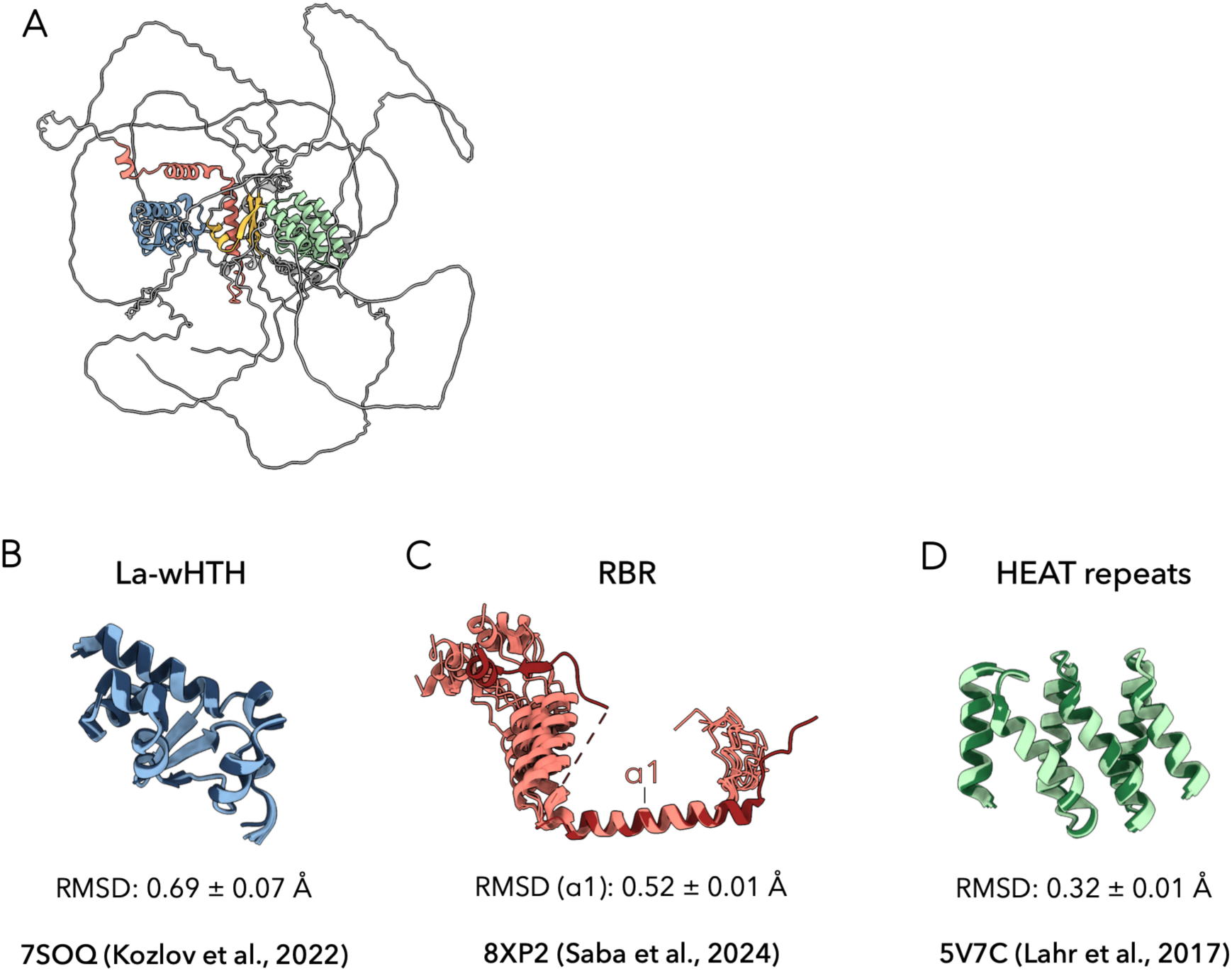
Comparison of AF3 models of full length LARP1 to solved structures. **A)** Representative AF3 model of full length LARP1. La-wHTH domain is depicted in blue, RRM in yellow with the RBR component in red, and HEAT repeats in green. **B)** Ten AF3 models (five each from two independent seeds) of full length LARP1 superimposed on the model of the La-wHTH domain from the solved structure reported in ^22^ (PDB 7SOQ). AF3 models are shown in light blue and the model from the solved structure is shown in dark blue. Only the La-wHTH domain (residues 400-487) is shown for clarity. Mean RMSD ± one standard deviation for residues 400-487 of the ten AF3 models compared to 7SOQ is reported. **C)** Ten AF3 models (five each from two independent seeds) of full length LARP1 superimposed on the model of the RBR domain from the solved structure reported in^14^ (PDB 8XP2). AF3 models are shown in light red and the model from the solved structure is shown in dark red. Only the RBR domain (residues 660-724) is shown for clarity. Mean RMSD ± one standard deviation for α1 (residues 673-691) of the ten AF3 models compared to 8XP2 is reported. The mean RMSD across the entirety of the RBR (residues 660-694; 708-724) is 15.83 ± 2.26 Å. **D)** Ten AF3 models (five each from two independent seeds) of full length LARP1 superimposed on the model of the HEAT repeat domain from the solved structure reported in^11^ (PDB 5V7C). AF3 models are shown in light green and the model from the solved structure is shown in dark green. Only the HEAT repeat domain (residues 890-999) is shown for clarity. Mean RMSD ± one standard deviation for residues 890-999 of the ten AF3 models compared to 5V7C is reported.

**Supplemental Figure S5.**
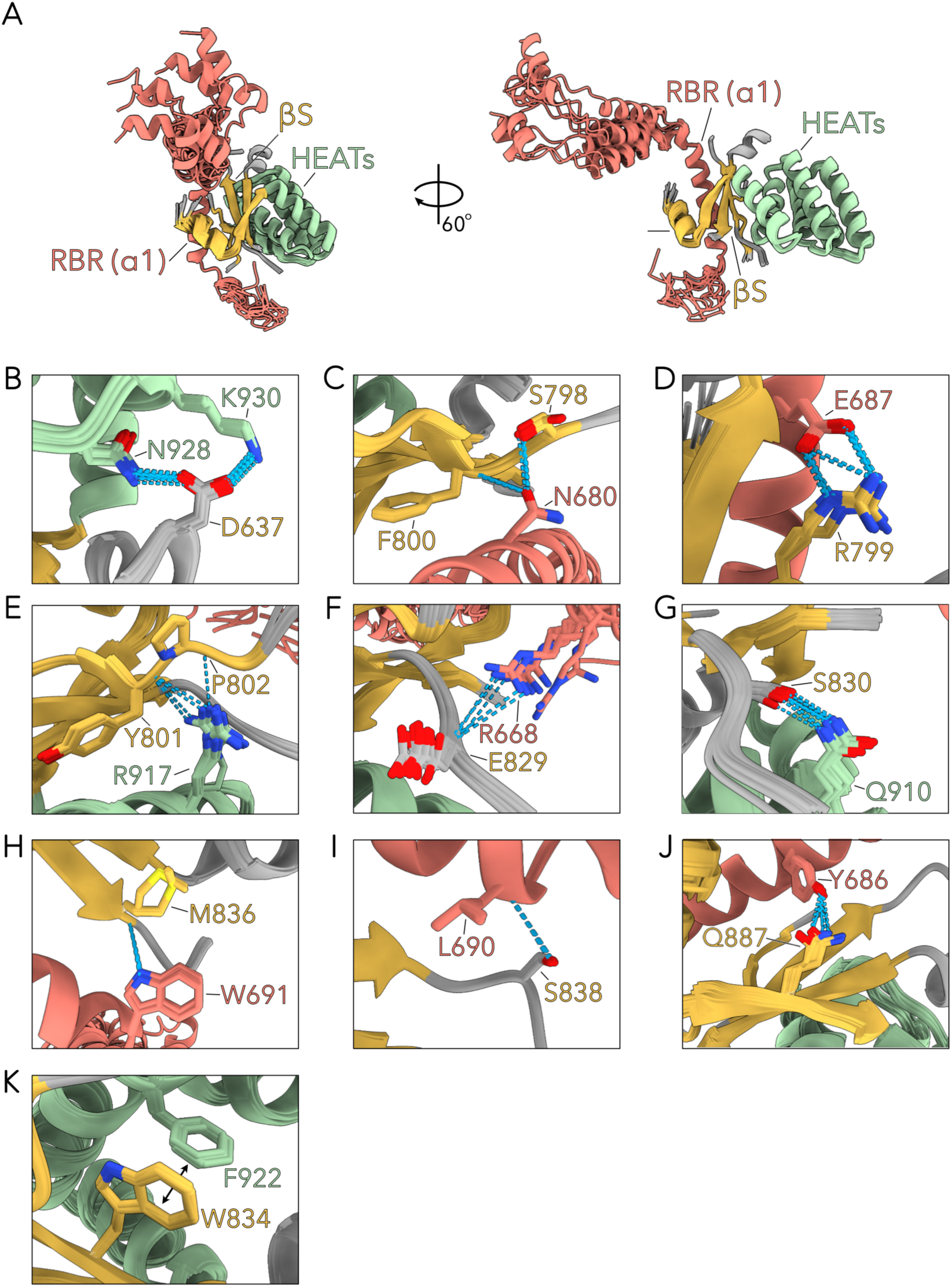
Structural analysis of LARP1 *in silico* predictions. **A)** Overlay of ten AF3 models (five each from two independent seeds) of full length LARP1, aligned by the RBR-RRM ɑ1. The RBR-RRM (yellow) with RBR component (red) and HEAT repeats (green) are shown. Residues 636-649, 660-724, 797-804, 828-839, and 874-999, encompassing the structured components of the RBR-RRM and HEAT repeats are shown for clarity. **B-J)** Zoom-ins depicting hydrogen bonding interactions between the RBR-RRM and HEAT repeats are shown for D637 with N928/K930 (**B**); S798/F800 with N680 (**C**); R799 with E687 (**D**); Y801/P802 with R917 (**E**); E829 with R668 (**F**); S830 with Q910 (**G**); M836 with W691 (**H**); S838 with L690 (**I**) and Q887 with Y686 (**J**). Hydrogen bonds are depicted as blue dashed lines. **K)** Zoom-in showing pi-stacking interaction between W834 and F922.

**Supplemental Figure S6.**
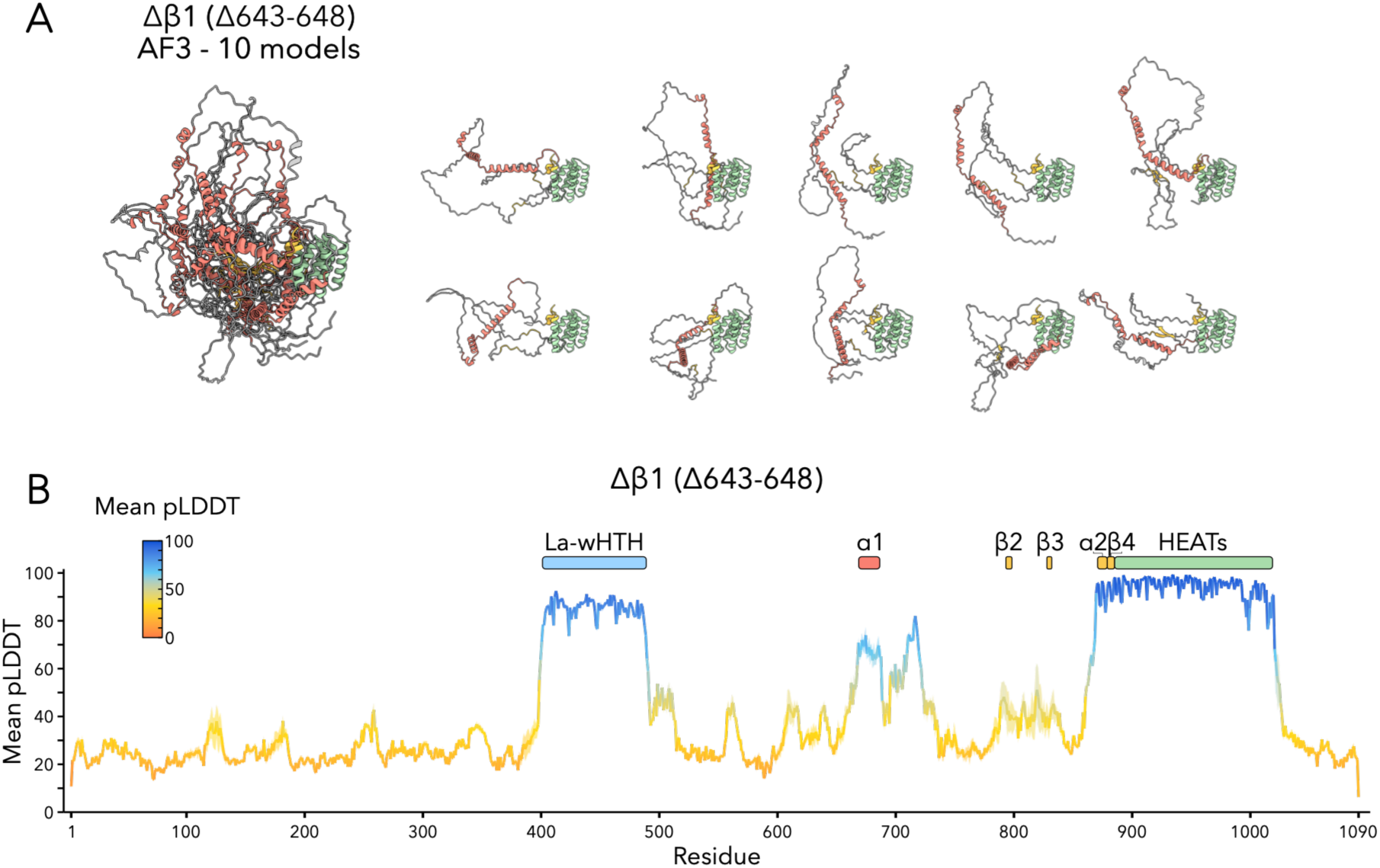
*In silico* deletion of the *β*1 strand of the RBR-RRM. **A)** Ten AF3 models (five models from each of two independent seeds) of Δ*β*1-LARP1 (Δ643-648) overlaid (left) and split into individual models (right). Models are aligned to the HEAT repeats (890-999). The region spanning the RBR-RRM to HEAT repeats (640-999) is shown for clarity. **B)** Mean per-residue pLDDT plot for Δ*β*1-LARP1 (Δ643-648) from ten AF3 predictions (five models generated from each of two independent seeds). The line plot represents the mean pLDDT scores and the shaded ribbon represents ± one standard deviation. Low pLDDT scores (red to green) indicate low confidence in the local folding of the α1, *β*2, and *β*3 strands of the RBR-RRM.

**Supplemental Figure S7.**
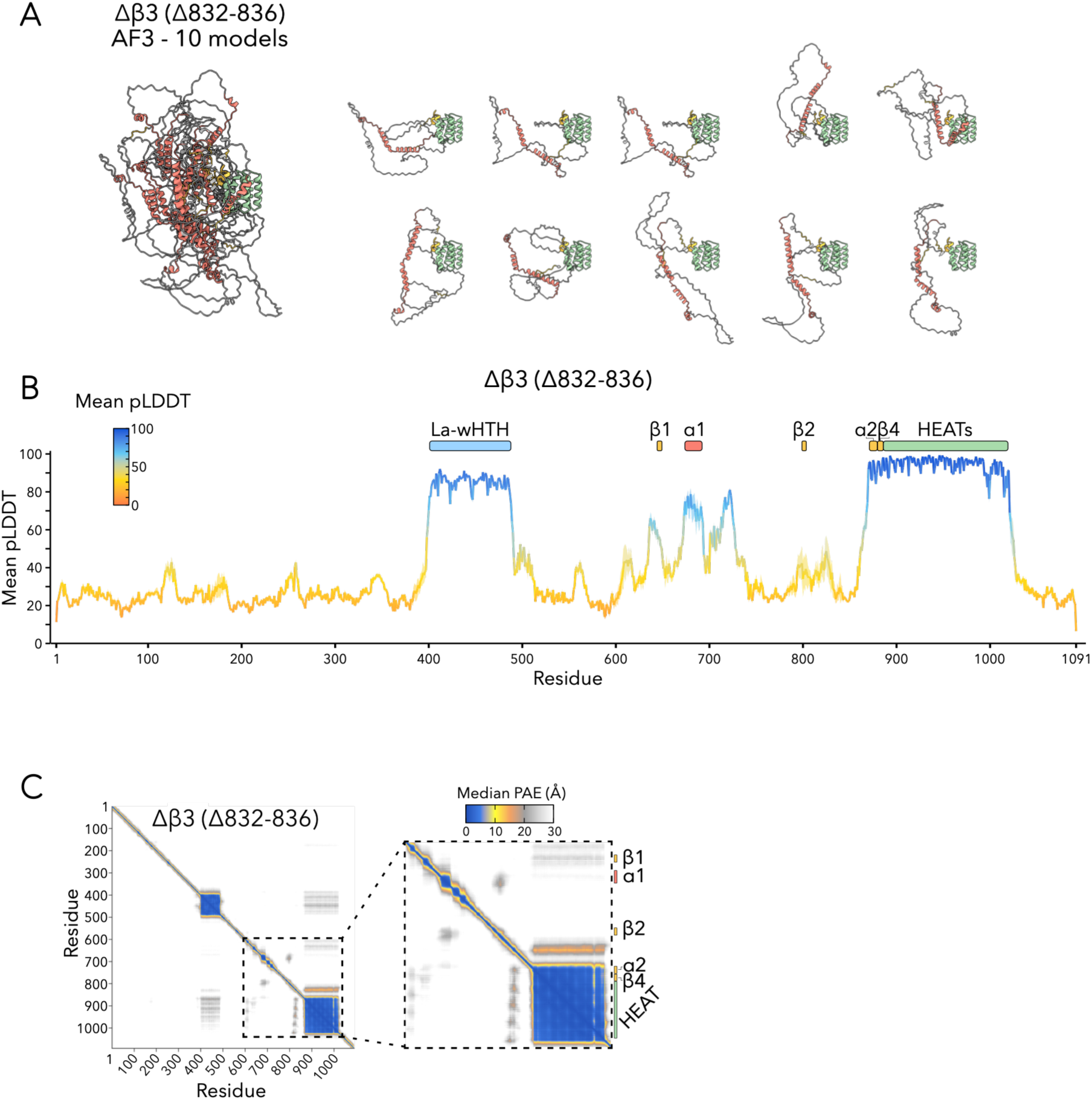
*In silico* deletion of the *β*3 strand of the RBR-RRM. **A)** Ten AF3 models (five models from each of two independent seeds) of Δ*β*3-LARP1 (Δ832-836) overlaid (left) and split into individual models (right). Models are aligned to the HEAT repeats (890-999). The region spanning the RBR-RRM to HEAT repeats (640-999) is shown for clarity. **B)** Mean per-residue pLDDT plot for Δ*β*3-LARP1 (Δ832-836) from ten AF3 predictions (five models generated from each of two independent seeds). The line plot represents the mean pLDDT scores and the shaded ribbon represents ± one standard deviation. Low pLDDT scores (red to green) indicate low confidence in the local folding of the *β*1, α1, and *β*2 strands of the RBR-RRM. **C)** Median predicted aligned error (PAE) plot for Δ*β*3-LARP1 (Δ832-836) from ten AF3 predictions (five models generated from each of two independent seeds). Zoom in of the region spanning the RBR-RRM and HEAT repeats is shown to the right. High PAE values (grey to white) demonstrate that the *β*S is not predicted to form and that the 40S-binding ɑ1 is not predicted to associate with the HEAT repeats.

**Supplemental Figure S8.**
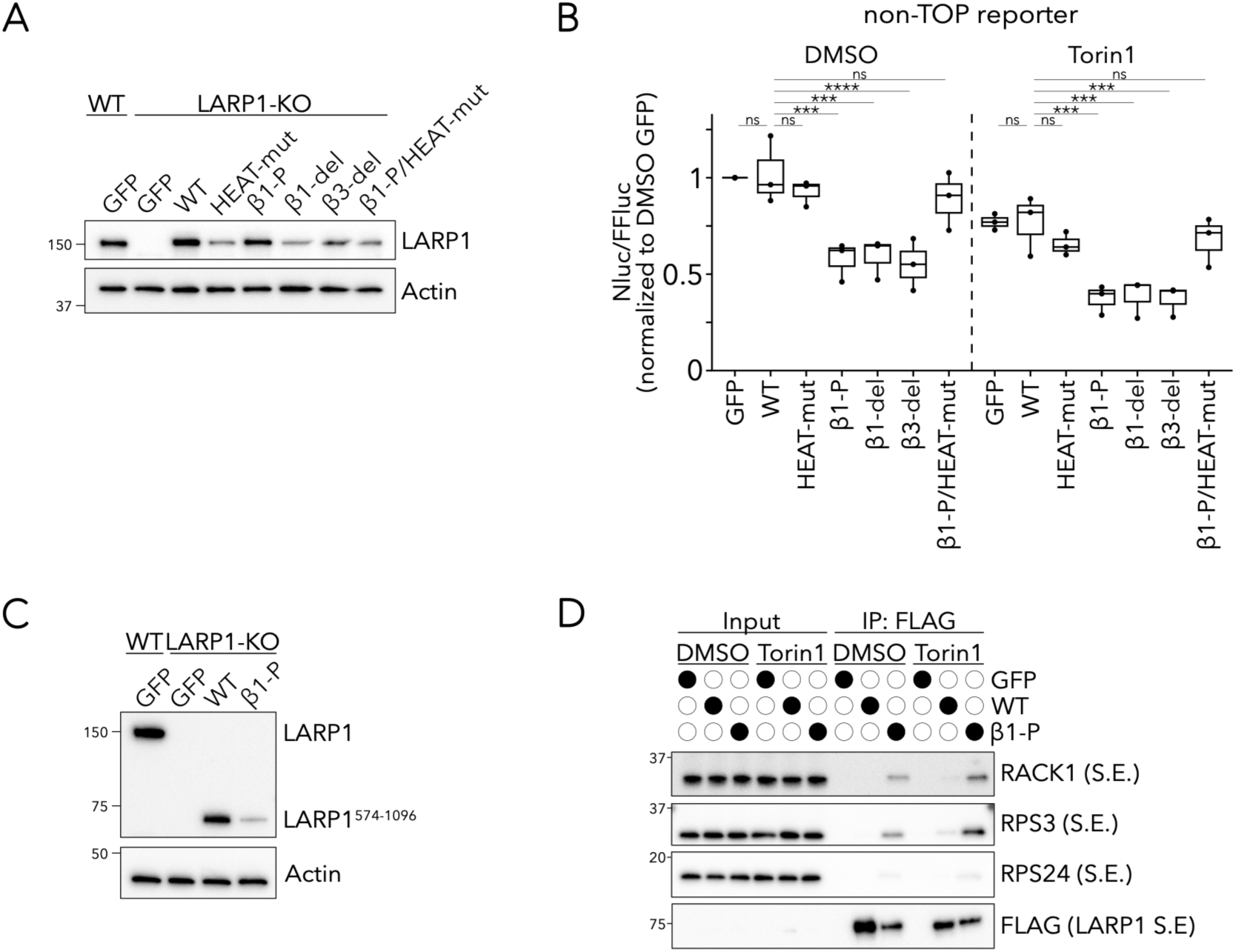
Characterization of LARP1 construct expression and non-TOP reporter activity. Related to Figure 2. **A)** Immunoblots comparing levels of endogenous LARP1 to levels of the indicated transfected LARP1 construct (or GFP) in LARP1-KO cells. These LARP1 expression levels apply to the TOP and non-TOP reporter assays in Figure 2B and Supplemental Figure S8B. Data is representative of two independent biological replicates. **B)** Expression of the non-TOP (UBL5-nLuc-PEST) reporter^14^. LARP1-KO cells were transfected with plasmids expressing the indicated LARP1 construct (or GFP) and treated with DMSO or 300 nM Torin1 for 1 hour. NLuc expression was normalized to FFLuc expressed on the same plasmid. Data points represent the mean of three technical replicates from three independent biological replicates. Whiskers represent minima and maxima; bounds of box represent the quartiles (25th and 75th percentile); center line represents the median. Values are normalized to the GFP-transfected, DMSO-treated sample. Two-way ANOVA with Dunnett’s correction for multiple comparisons; n.s. not significant, *** p ≤ 0.001. **C)** Immunoblots comparing levels of endogenous LARP1 to levels of the indicated transfected LARP1^574-1096^ construct (or GFP) in LARP1-KO cells. These truncated LARP1 expression levels apply to the ribosome binding assays in Figure 2C and Supplemental Figure S8D. Data is representative of two independent biological replicates. **D)** Short exposures (S.E.) for RACK1, RPS3, RPS24, and FLAG (LARP1) immunoblots shown in Figure 2C.

**Supplemental Figure S9.**
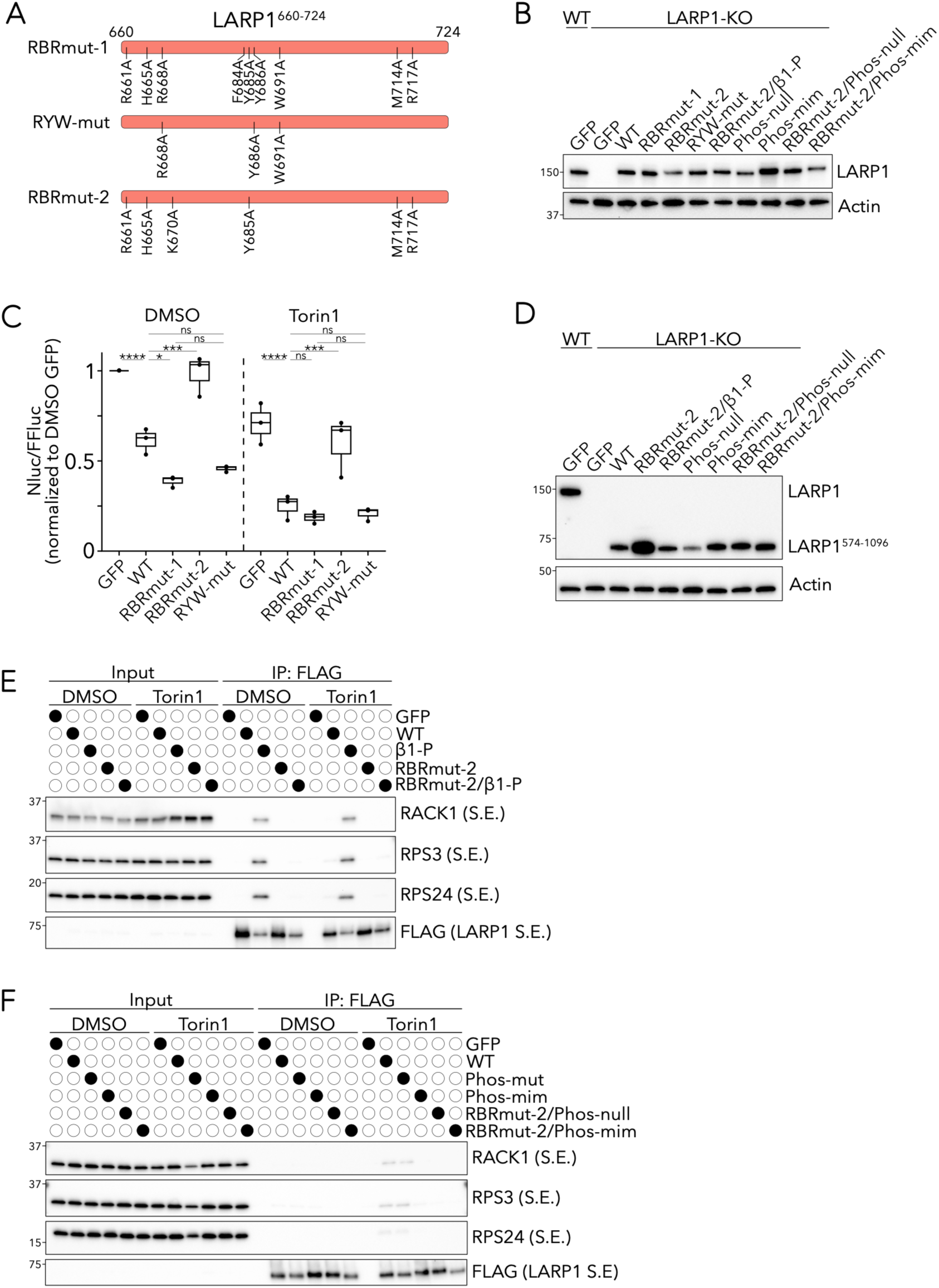
RBRmut-2 LARP1 is deficient for 40S binding and TOP repression. **A)** Schematic depicting the LARP1 RBR (residues 660-724) with the indicated point mutations used in RBRmut-1 (top), RYW-mut (middle), and RBRmut-2 (bottom) LARP1. **B)** Immunoblots comparing levels of endogenous LARP1 to levels of the indicated transfected LARP1 construct (or GFP) in LARP1-KO cells. These LARP1 expression levels apply to the TOP reporter assays in Figures 2E-F and Supplemental Figure S9C. Data is representative of two independent biological replicates. **C)** Expression of the TOP (RPL39-nLuc-PEST) reporter^14^. LARP1-KO cells were transfected with plasmids expressing the indicated LARP1 construct (or GFP) and treated with DMSO or 300 nM Torin1 for 1 hour. NLuc expression was normalized to FFLuc expressed on the same plasmid. Data points represent the mean of three technical replicates from three independent biological replicates. Whiskers represent minima and maxima; bounds of box represent the quartiles (25th and 75th percentile); center line represents the median. Values are normalized to the GFP-transfected, DMSO-treated sample. Two-way ANOVA with Tukey’s correction for multiple comparisons; n.s. not significant, * p ≤ 0.05, *** p ≤ 0.001, **** p ≤ 0.0001. **D)** Immunoblots comparing levels of endogenous LARP1 to levels of the indicated transfected LARP1^574-1096^ construct (or GFP) in LARP1-KO cells. These truncated LARP1 expression levels apply to the ribosome binding assays in Figures 2D and 2G and Supplemental Figures S9E-F. Data is representative of two independent biological replicates. **E)** Short exposures (S.E.) for RACK1, RPS3, RPS24, and FLAG (LARP1) immunoblots shown in Figure 2D. **F)** Short exposures (S.E.) for RACK1, RPS3, RPS24, and FLAG (LARP1) immunoblots shown in Figure 2G.

**Supplemental Figure S10.**
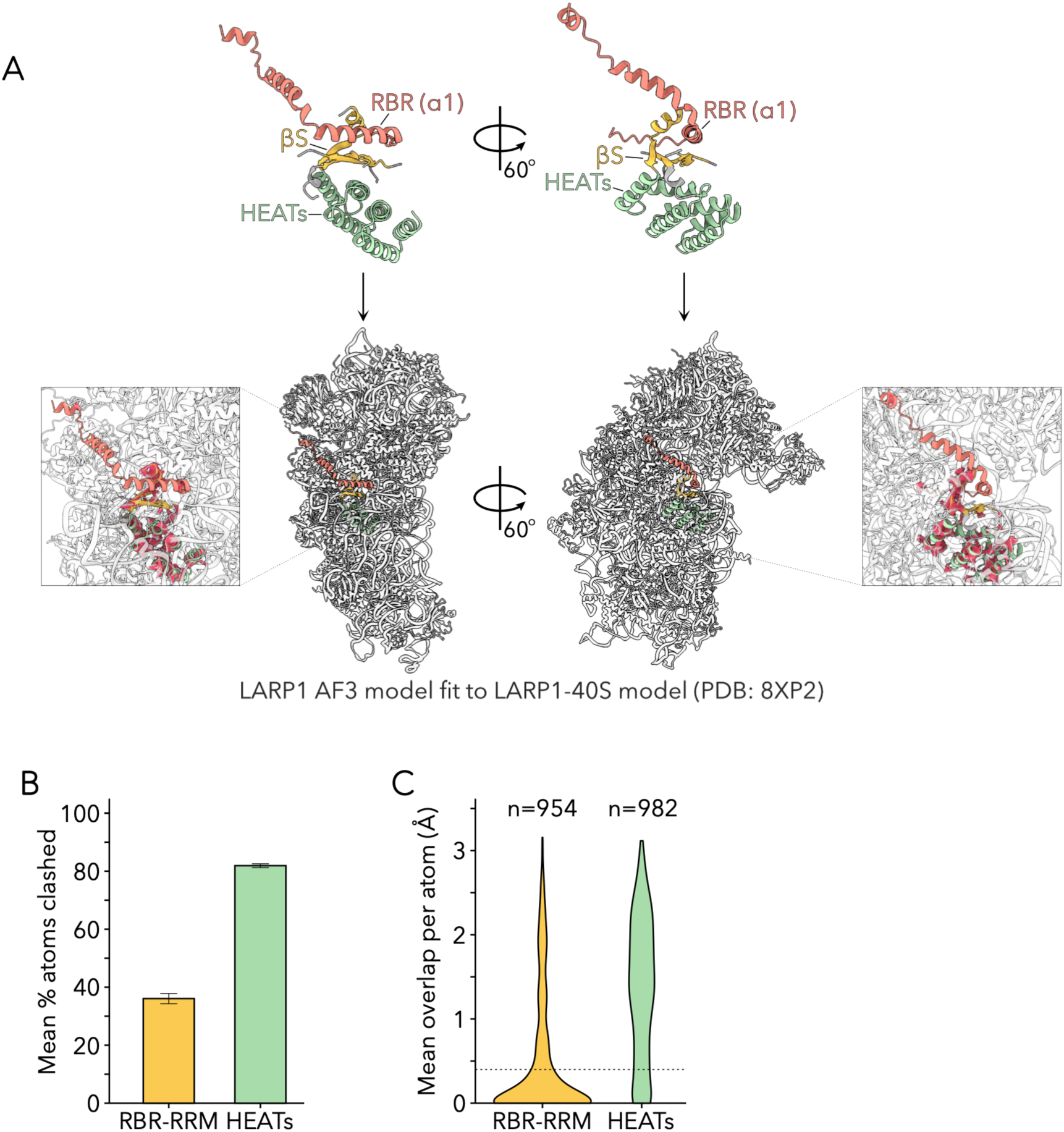
40S binding to the ɑ1 helix is not compatible with the predicted RBR-RRM/HEAT scaffold. **A)** Representative model of the ɑ1 helix (residues 673-691) from the AF3-predicted structure of WT LARP1 fit to the solved structure of the same helix bound to the mRNA channel of the 40S subunit^14^ (8XP2; residues 673-691; mean RMSD from ten AF3 models = 0.52 Å). The AF3-predicted RRM and HEAT repeats are buried within the 40S subunit, demonstrating that 40S binding to the ɑ1 helix is incompatible with the predicted structure of the RBR-RRM/HEAT scaffold. Clashes (overlap > 0.4 Å^72^) between atoms of predicted RBR-RRM/HEAT scaffold and the 40S subunit are shown in red within the inset. LARP1 residues 636-649, 660-724, 797-804, 828-839, and 874-999, encompassing the structured components of the RBR-RRM (yellow, with RBR component in red) and HEAT repeats (green) are shown for clarity. **B)** Mean percent of atoms across ten AF3 models (five models from each of two independent seeds) in the predicted RBR-RRM/HEAT scaffold that clash with atoms in the 40S subunit (overlap > 0.4 Å^72^). Error bars represent ± one standard deviation. Only atoms within the predicted structured components of the RBR-RRM (residues 636-649, 660-724, 797-804, 828-839, and 874-999) are included in the analysis. **C)** Mean overlap between an atom in the RBR-RRM or HEAT repeat domains and an atom in the 40S subunit across ten AF3 models (five models from each of two independent seeds). The dashed line represents an overlap of 0.4 Å; all values greater are considered a serious clash^72^. The majority of atoms in the HEAT repeats and many atoms in the RBR-RRM exhibit clashes over 0.4 Å. If an atom in LARP1 clashed with multiple atoms in the 40S subunit, only the clash with the greatest overlap is included. Only atoms within the predicted structured components of the RBR-RRM (residues 636-649, 660-724, 797-804, 828-839, and 874-999) are included in the analysis.

**Supplemental Figure S11.**
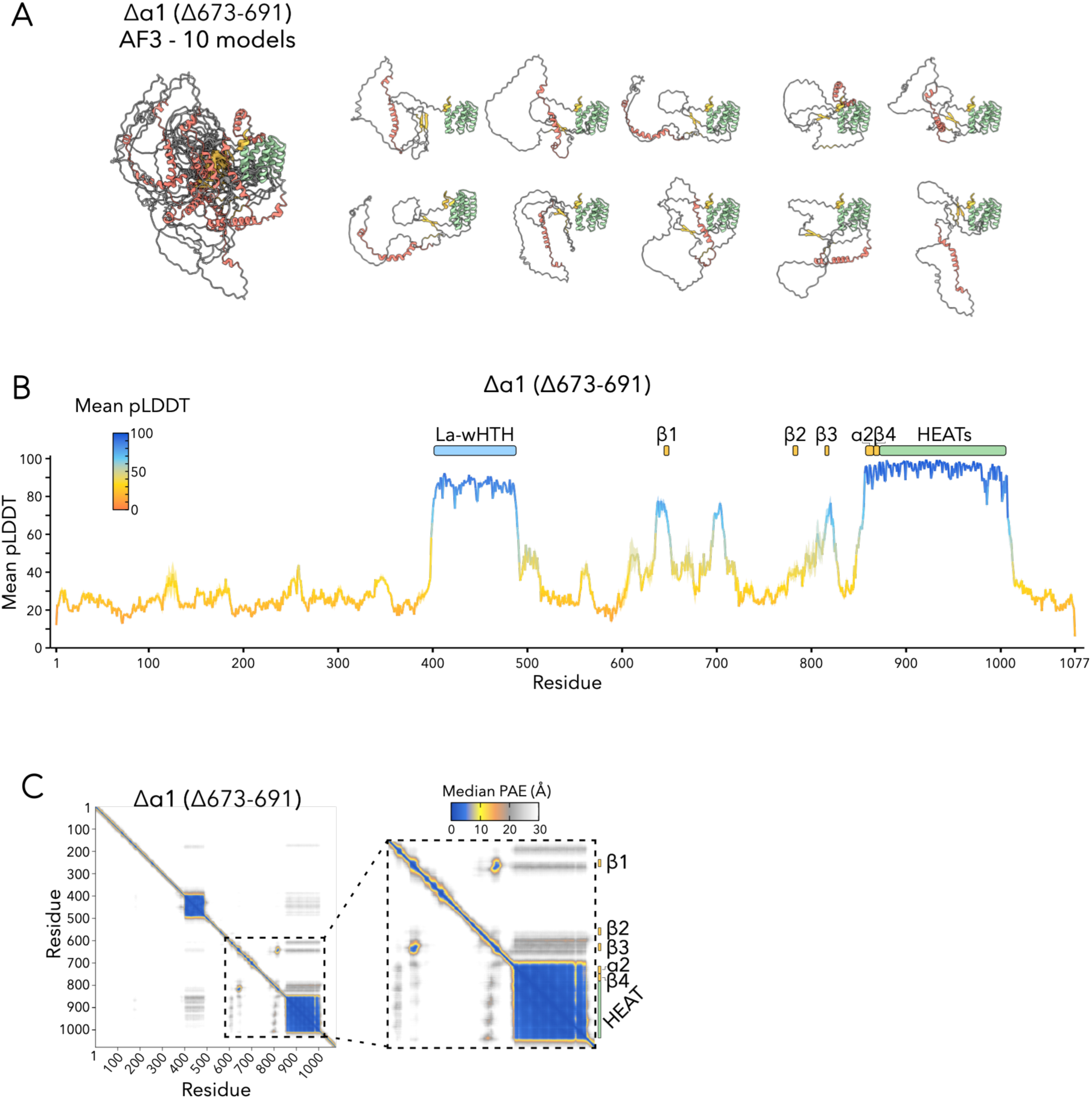
*In silico* deletion of the ɑ1 helix of the RBR-RRM. **A)** Ten AF3 models (five models from each of two independent seeds) of Δɑ1-LARP1 (Δ673-691) overlaid (left) and split into individual models (right). Models are aligned to the HEAT repeats (890-999). The region spanning the RBR-RRM to HEAT repeats (640-999) is shown for clarity. **B)** Mean per-residue pLDDT plot for Δɑ1-LARP1 (Δ673-691) from ten AF3 predictions (five models generated from each of two independent seeds). The line plot represents the mean pLDDT scores and the shaded ribbon represents ± one standard deviation. **C)** Median predicted aligned error (PAE) plot for Δɑ1-LARP1 (Δ673-691) from ten AF3 predictions (five models generated from each of two independent seeds). Zoom in of the region spanning the RBR-RRM and HEAT repeats is shown to the right. High PAE values (grey to white) demonstrate that the HEAT repeats are displaced from the core *β*1 and *β*3 strands.

**Supplemental Figure S12.**
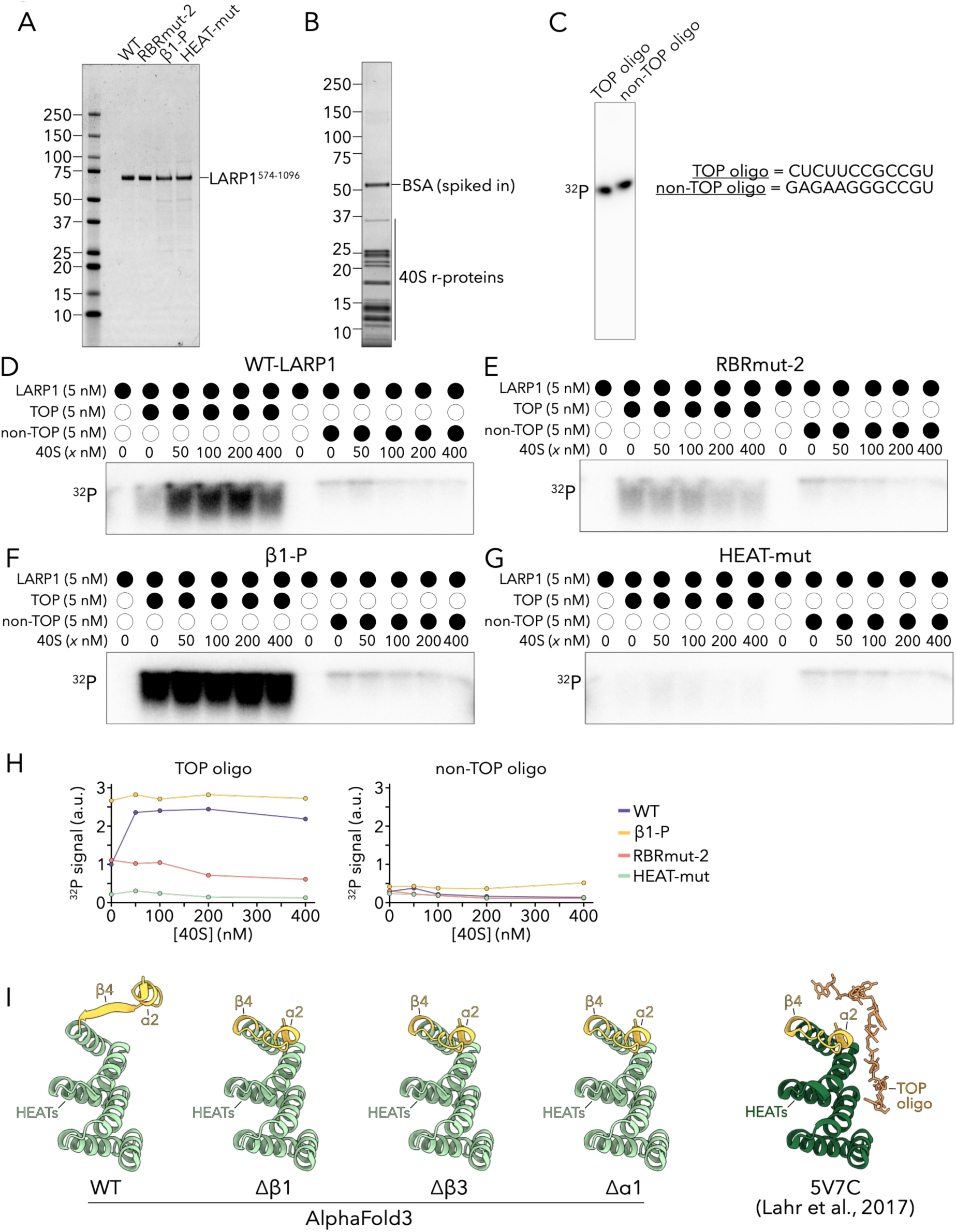
LARP1 couples 40S binding and TOP binding *in vitro*. Related to Figure 3. **A)** Coomassie-stained gel demonstrating WT and mutant LARP1^574-1096^ purifications. **B)** Coomassie-stained gel demonstrating 40S subunit purification. **C)** ^32^P exposure demonstrating capped TOP and non-TOP oligonucleotide purifications and equal loading (5 nM) used for all experiments in Figure 3 and Supplemental Figure S12. **D**-**G)** Identical setup to Figures 3B-E using higher concentrations of 40S subunits. Panels (**D**-**G**) were performed in a single experiment; levels of bound oligonucleotide can be directly compared between panels. **H)** Quantification of ^32^P signal intensity in (**D**-**G**) (a.u. arbitrary units). Values are normalized to WT LARP1 incubated without 40S subunits. **I)** Representative AF3 models of the RBR-RRM ɑ2 helix, *β*4 strand, and HEAT repeats in WT-LARP1 compared to Δ*β*1 (Δ643-648), Δ*β*3 (Δ832-836), Δɑ1 (Δ673-691), or the solved structure of this region bound to a TOP oligo (PDB 5V7C)^11^. Mutant LARP1 models exhibit a predict refolding of *β*4 to a helical conformation identical to that observed in the solved structure.LARP1 residues 875-999 are shown. Mean RMSD ± one standard deviation for residues 875-999 is calculated from ten AF3 predictions (five models generated from each of two independent seeds) compared to the solved structure (RMSDs: WT 5.32 ± 0.04 Å, Δ*β*1 0.36 ± 0.02 Å, Δ*β*3 0.36 ± 0.01 Å, Δɑ1 0.36 ± 0.01 Å). For clarity, a single AF3 model is reproduced from Supplemental Figures S1A (WT), S6A (Δ*β*1), S7A (Δ*β*3), and S11A (Δɑ1).

**Supplemental Figure S13.**
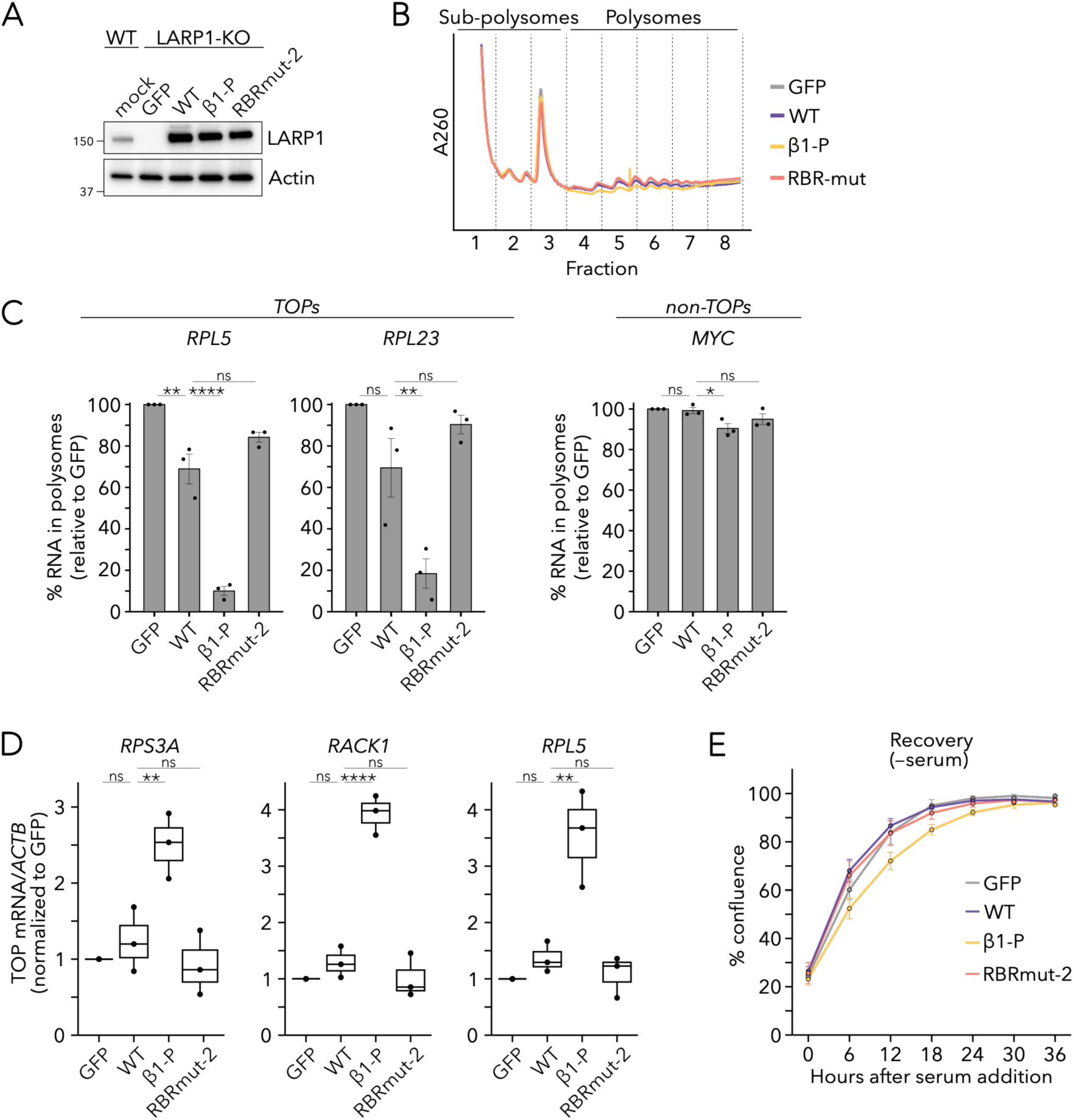
The predicted RBR-RRM/HEAT scaffold regulates endogenous TOPs. Related to Figure 4. **A)** Immunoblots comparing levels of endogenous LARP1 to levels of the indicated transfected LARP1 construct (or GFP) in LARP1-KO cells for the assays shown in Figures 4A-B and Supplemental Figures S13B-C. Data is representative of three independent biological replicates. **B)** Representative A260 absorbance trace of sucrose gradient samples used in Figures 4A-B and Supplemental Figure S13C. Fractions are labeled below the trace. Fractions 1-3 and 4-8 were pooled as sub-polysomes and polysomes, respectively. Data is representative of three independent biological replicates. **C)** Distributions of additional TOPs and non-TOPs from experiment shown in Figures 4A-B. Bars represent the mean of three independent biological replicates. Data points represent the mean of three technical replicates from three independent biological replicates and error bars represent the SEM. Values are normalized to the GFP-transfected sample. One-way ANOVA with Dunnett’s correction for multiple comparisons; n.s. not significant, * p ≤ 0.05, ** p ≤ 0.01, **** p ≤ 0.0001. **D)** Steady-state TOP mRNA levels measured by RT-qPCR relative to *ACTB* mRNA. LARP1-KO cells were transfected with the indicated LARP1 construct (or GFP) for 48 hours. Data points represent the mean of three technical replicates for three independent biological replicates. Whiskers represent minima and maxima; bounds of box represent the quartiles (25th and 75th percentile); center line represents the median. Values are normalized to the GFP-transfected sample. One-way ANOVA with Dunnett’s correction for multiple comparisons; n.s. not significant, ** p ≤ 0.01, **** p ≤ 0.0001. **E)** Growth curves over 36 hours following 24 hours of serum starvation of LARP1-KO cells transfected with the indicated LARP1 construct (or GFP) at endogenous levels. Data points represent the mean of three technical replicates. Data is representative of three independent biological replicates. Error bars represent one standard deviation from the mean across technical replicates.

**Supplemental Table 1.** LARP1 representative domain architectures and boundaries across eukaryotic phylogenies.

